# Mathematical modeling and biochemical analysis support partially ordered CaM-MLCK binding

**DOI:** 10.1101/2022.06.08.495195

**Authors:** Melissa JS MacEwen, Domnita-Valeria Rusnac, Henok Ermias, Timothy M Locke, Hayden E Gizinski, Joseph P Dexter, Yasemin Sancak

## Abstract

Activation of myosin light-chain kinase (MLCK) by calcium ions (Ca^2+^) and calmodulin (CaM) plays an important role in numerous cellular functions including vascular smooth muscle contraction and cellular motility. Despite extensive biochemical analysis of this system, aspects of the mechanism of activation remain controversial, and competing theoretical models have been proposed for the binding of Ca^2+^ and CaM to MLCK. The models are analytically solvable for an equilibrium steady state and give rise to distinct predictions that hold regardless of the numerical values assigned to parameters. These predictions form the basis of a recently proposed, multi-part experimental strategy for model discrimination. Here we implement this strategy by measuring CaM-MLCK binding using an *in vitro* FRET system. This system uses the CaM-binding region of smooth muscle MLCK protein to link two fluorophores to form an MLCK FRET Reporter (FR). Biochemical and biophysical experiments have established that FR can be reliably used to analyze MLCK-CaM binding. We assessed the binding of either wild-type CaM, or mutant CaM with one or more defective EF-hand domains, to FR. Interpretation of binding data in light of the mathematical models suggests a partially ordered mechanism for binding of CaM to MLCK. Complementary data collected using orthogonal approaches that directly quantify CaM-MLCK binding further supports our conclusions.

## Introduction

Calmodulin (CaM) is a calcium-binding protein found ubiquitously in the cytosol of eukaryotic cells. CaM contains two globular domains joined by a flexible linker. The N-terminus and C-terminus domains each have a pair of EF-hand motifs and are each capable of binding two calcium ions (Ca^2+^) (*1*). CaM acts as a regulator or effector in numerous cellular processes, ranging from muscle contraction to glycogen metabolism and synaptic plasticity (*2-4*). The Ca^2+^-mediated binding of CaM and either smooth muscle myosin light chain kinase or nonmuscle myosin light chain kinase (henceforth collectively referred to as “MLCK”) is central to functions such as vascular smooth muscle contraction and cell motility. CaM-MLCK binding takes place as follows: Ca^2+^ enters the cytosol and binds to the four Ca^2+^ binding sites of CaM, leading to a dramatic change in CaM protein conformation (*5*). CaM then binds to the CaM-binding domain of MLCK, triggering a conformational change in MLCK that activates the kinase by displacing an autoinhibitory sequence from the kinase’s catalytic domain (*6, 7*). Finally, activated MLCK phosphorylates the 20-kDa regulatory light chains of myosin II, resulting in contraction caused by myosin cross-bridges moving along actin filaments.

Multiple formal models have been proposed for the activation of MLCK by Ca^2+^ and CaM (*8-11*). Given the cooperative nature of Ca^2+^ binding at both the C- and N-terminus of CaM, each of these models treats CaM as having two Ca^2+^ binding sites. The model of Brown *et al*. (*8*) and Fajmut, Brumen *et al*. (*9*) makes no further assumptions, giving rise to an eight-state reaction network in which MLCK may bind to CaM before or after Ca^2+^ (“Model 1” in **Fig. 1A**). Other previously proposed models are truncations of this network, in which the binding of Ca^2+^ and MLCK is either partially (*10*) or fully (*11*) ordered (“Model 2” and “Model 3” in **Fig. 1A**, respectively). Accordingly, Model 1 corresponds to a fully random binding mechanism (Ca^2+^ and MLCK can bind to CaM in any order), Model 2 to a partially ordered mechanism (MLCK can bind to CaM after Ca^2+^ is bound at the C-terminus), and Model 3 to a fully ordered mechanism (MLCK can bind to CaM only after Ca^2+^ is bound at both the C-terminus and N-terminus).

**Figure 1:**
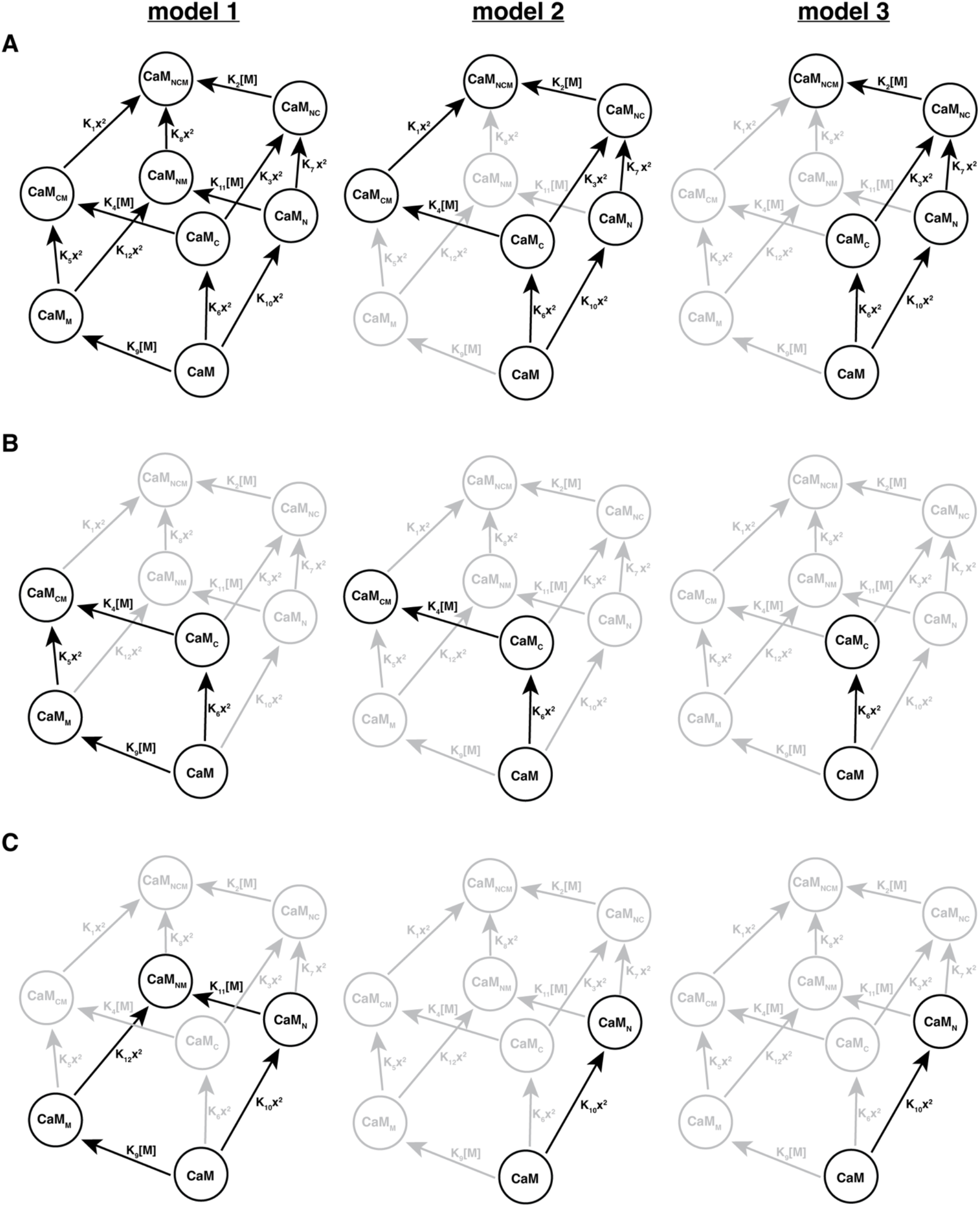
Network diagrams for the binding of Ca^2+^ and MLCK to CaM. Model 1 is the unordered model proposed by Brown *et al*. (*8*) and Fajmut, Brumen *et al*. (*9*); models 2 and 3 correspond to the partially and fully ordered models of Fajmut, Jagodic *et al*. (*10*) and Kato *et al*. (*11*), respectively. **(A)** shows the networks for CaM^WT^, **(B)** for CaM^21A,57A^, and **(C)** for CaM^94A,130A^ Models 2 and 3 are truncations of Model 1; omitted edges and vertices are shown in gray. In each network *x* denotes [Ca^2+^], and *M* denotes MLCK. The subscripts N and C indicate Ca^2+^ binding at the N-terminus and C-terminus EF hands of CaM, respectively, and the subscript M indicates MLCK binding. The figure is adapted from Dexter *et al*. (*12*).

In recent work, Dexter *et al*. showed that this class of models is analytically solvable for a CaM/MLCK system at both thermodynamic equilibrium and steady state, and that each model predicts distinct steady-state behavior in certain Ca^2+^ concentration regimes (*12*). Importantly, these predictions hold regardless of the numerical values assigned to the model parameters (i.e., the equilibrium constants in **Fig. 1A**), allowing us to develop a strategy for model discrimination that does not require extensive parameter estimation.

Our biochemical approach to testing the predictions of the models centers on the use of an established fluorescence resonance energy transfer (FRET) reporter that we term FR. FR has been employed to characterize CaM- and Ca^2+^-dependent activation of MCLK both *in vivo* and *in vitro* (*13-19*). FR acts as a stand-in for the wild-type MLCK protein; it is composed of an EYFP and ECFP FRET pair, which are linked by the CaM-binding domain of smooth muscle MLCK (*14*). Binding of CaM to FR interferes with FRET, enabling analysis of FR-CaM binding in the form of the ratio of EYFP and ECFP emission (F480/F535) after excitation at 430 nm (**Fig. 2A**). For our FRET-based binding assays, we measure FR-CaM binding across a wide range of [Ca^2+^] for wild-type CaM (CaM^WT^) and three different CaM mutants: CaM with its N-terminus EF-hands mutated (D21A and D57A mutations, CaM^21A,57A^; **Fig. 1B**), CaM with its C-terminus EF-hands mutated (D94A and D130A mutations, CaM^94A,130A^; **Fig. 1C**) and CaM with all of its EF-hands mutated (D21A, D57A, D94A, D130A,CaM^21A,57A,94A,130A^). Each of the site-directed Asp to Ala mutations we performed has been demonstrated to prevent Ca^2+^ binding, and to have only a slight impact on the structure of each CaM EF hand (*20*).

**Figure 2:**
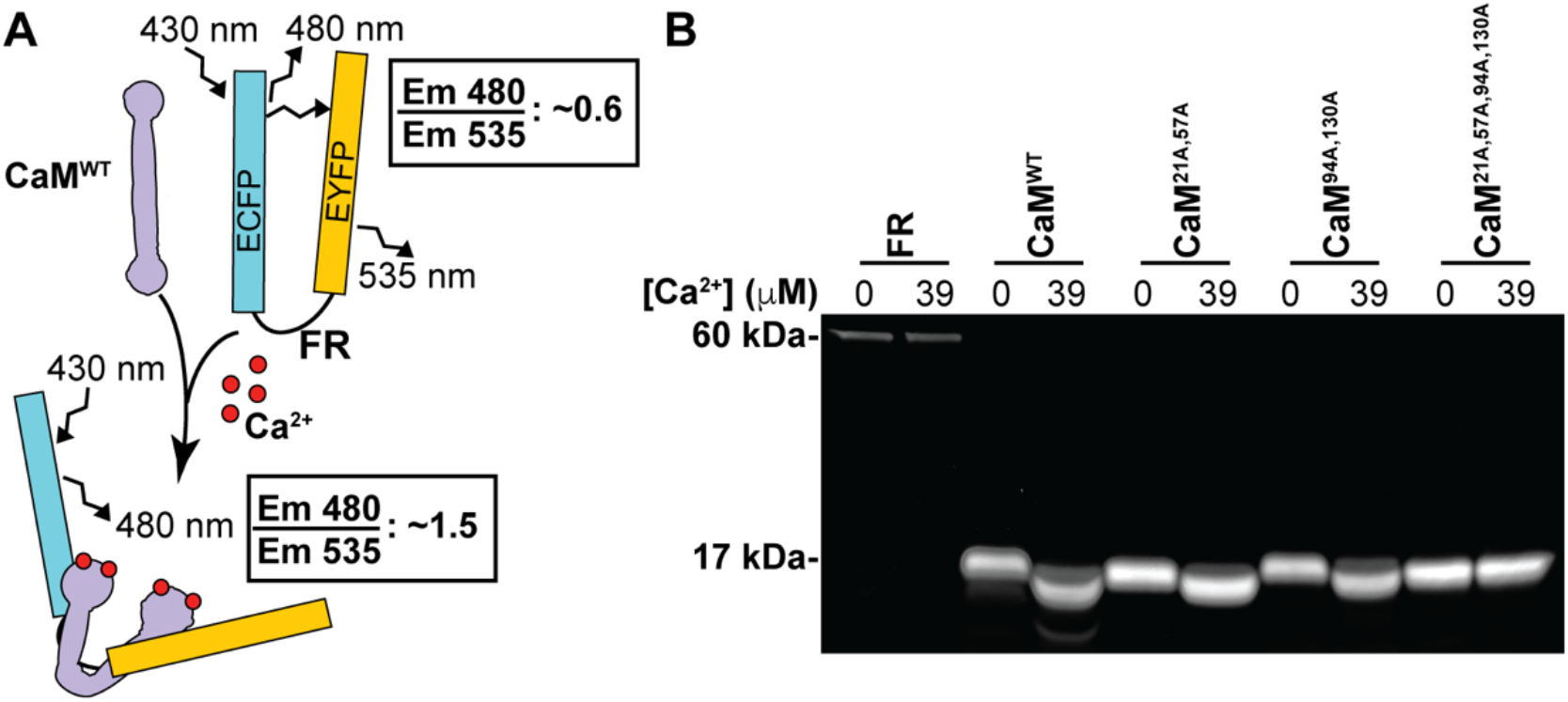
MLCK-based FRET assay reports MCLK-CaM binding. **(A)** Schematic showing assay principle. In the presence of Ca^2+^, CaM binds to the CaM-binding domain of FR, leading to a decrease in FRET and reduced emission at 535 nm. **(B)** Representative gel image of purified FR and CaM proteins visualized with SYPRO™ Ruby protein gel stain following SDS-PAGE. Samples were incubated in buffers containing either 0 μM Ca^2+^ or 39 μM free Ca^2+^ before SDS-PAGE. 8 μg of each CaM protein and 800 ng of FR was loaded.

We use a multi-step strategy to discriminate between these models. We first compare Model 1 to Models 2 and 3, and then we compare Model 3 to Models 1 and 2. During the first step, FR-CaM binding at zero [Ca^2+^] is determined as Model 1, but not Models 2 and 3, predicts binding under these conditions. In the second step, binding of FR to a CaM with impaired N-terminus Ca^2+^ (CaM^21A,57A^) is measured at high [Ca^2+^]. Models 1 and 2, but not Model 3, predict binding between FR and this mutant CaM at high [Ca^2+^]. Finally, we use a CaM mutant with impaired C-terminus Ca^2+^ binding (CaM^94A,130A^) to evaluate additional predictions of Model 2. Because this approach assesses binding behavior predicted by the three proposed models, no parameter estimation is required or employed.

Our binding measurements using the FR falsify key predictions of both Model 1 (random binding) and Model 3 (fully ordered binding) but are consistent with multiple distinct predictions of Model 2, which strongly suggests that CaM-MLCK binding follows a partially ordered mechanism. We further validate our key findings and the utility of the FR system using orthogonal experimental techniques and assessing binding between full length MLCK-FLAG protein and CaM.

## Results

For all binding experiments, we purified either FLAG-tagged FR or MLCK from HEK 293T cells after transient transfection, and purified His-tagged CaM^WT^ and CaM mutants (CaM^21A,57A^, CaM^94A,130A^, and CaM^21A,57A,94A,130A^) after bacterial expression. Gel electrophoresis and SYPRO™ Ruby staining of purified proteins (**Fig. 2B**), as well as mass spectrometry analysis (**Fig. S1**), established that the protein preparations were free of major contaminants. It is well documented that the electrophoretic mobility of CaM in SDS-PAGE changes following Ca^2+^ binding-induced conformational change. This gel mobility shift is commonly used as a readout of Ca^2+^-CaM binding (*21-24*). We observed an expected Ca^2+^-dependent gel mobility shift in our CaM^WT^, CaM^21A,57A^, and CaM^94A,130A^ proteins (**Fig. 2B**)(*25*), but observed no gel mobility shift for FR or CaM^21A,57A,94A,130A^ in the presence of Ca^2+^. These data suggest that our purified CaM proteins each retain their expected Ca^2+^ -binding affinity.

Having validated our experimental tools, we collected FRET-based FR-CaM binding data to use in our model discrimination strategy. The previous theoretical analysis identified a two-part strategy for distinguishing between the three models of CaM-MLCK binding, which is described in detail in Dexter *et al*. (*12*). The analysis involves deriving algebraic expressions for the fraction of total MLCK (*F*) that is predicted to bind to CaM as a function of free [Ca^2+^] for each of the models. Plots of *F* are shown in **Fig. 3A** for CaM^WT^, CaM^21A,57A^, CaM^94A,130A^, and CaM^21A,57A,94A,130A^ (assuming the reference values for the equilibrium constants compiled by Fajmut, Brumen *et al*. (*9*) and the concentrations of CaM and MLCK used in our experiments). The model discrimination strategy rests on two predictions that differ between the three models and that hold true regardless of the specific numerical values assigned to the model parameters, such as the equilibrium constants in **Fig. 1**. The first is that only Model 1 predicts non-zero binding of MLCK in zero [Ca^2+^], with the fraction bound given by the following expression:

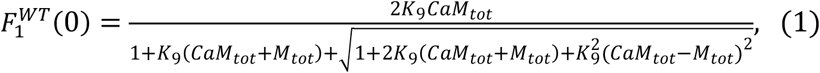

where CaM_tot_ and MLCK_tot_ denote total CaM and total MLCK, respectively. The second is that only Model 3 predicts zero binding of MLCK to a CaM mutant with nonfunctional N-terminal EF hands, such as CaM^21A,57A^, at any free [Ca^2+^]. In contrast, both Models 1 and 2 predict non-zero binding, with the fraction in the high-[Ca^2+^] limit given by

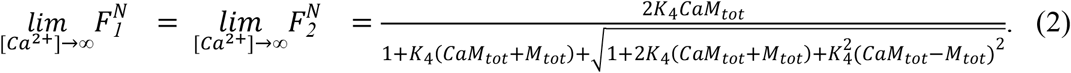

The algebraic structure of Eqs. 1 and 2 is identical; the only difference between the two expressions is which equilibrium constant appears (K_9_ in Eq. 1, K_4_ in Eq. 2). As discussed in Dexter *et al*. (*12*), this similarity is a consequence of the fact that the reaction network reduces to a bimolecular reaction in both the low- and high-[Ca^2+^] limits.

**Figure 3:**
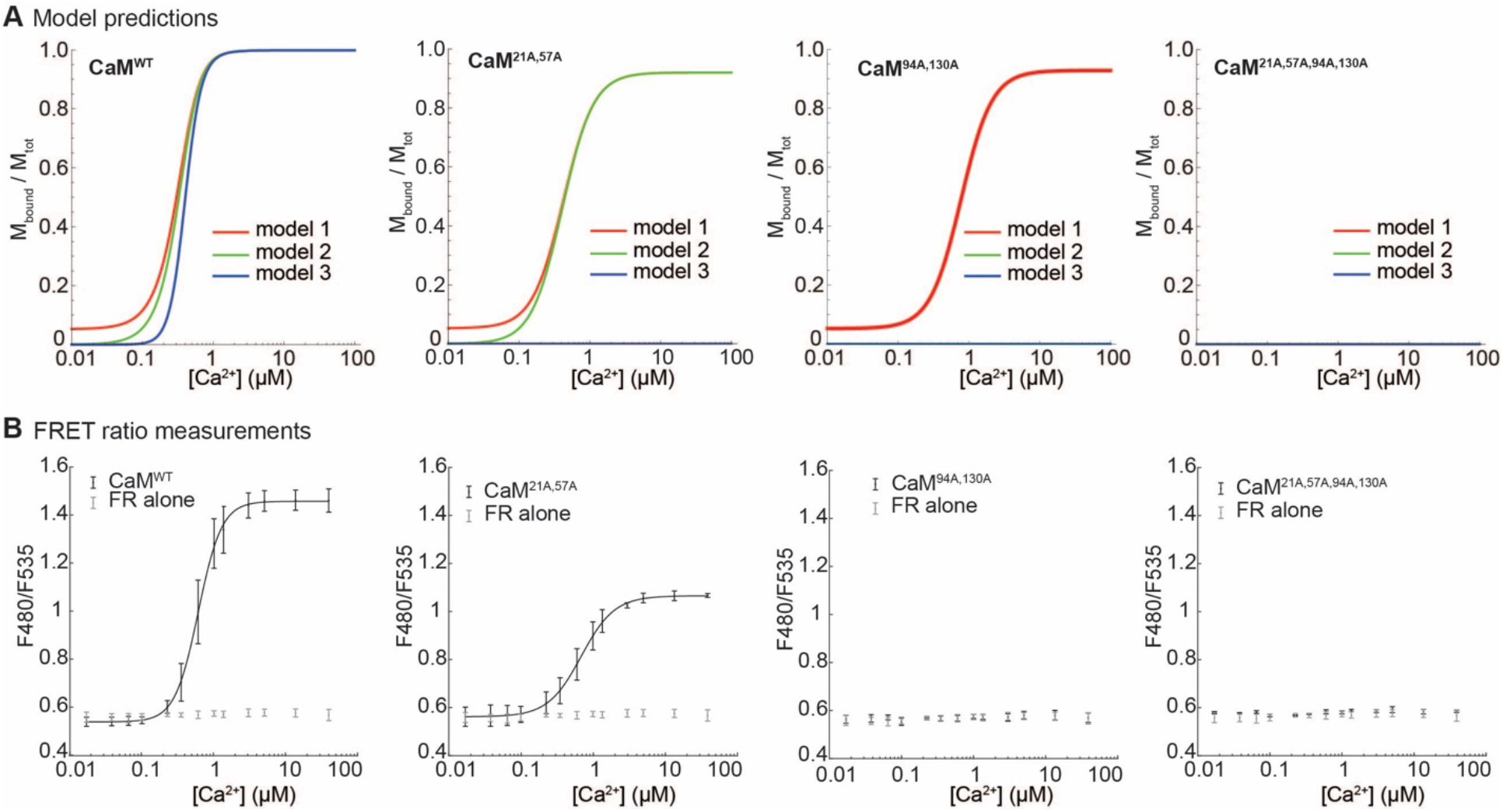
Predictions of mathematical models of MLCK binding and FRET measurements in the presence of excess CaM relative to FR. **(A)** Theoretical CaM-MLCK binding curves for CaM^WT^, CaM^21A,57A^, CaM^94A,130A^, and CaM^21A,57A,94A,130A^. The predicted curves were calculated assuming [CaM] = 0.713 μM and [MLCK] = 0.0237 μM (the concentrations of CaM and FR used in the main experiments) and the reference parameter values from Fajmut, Brumen *et al*. (*9*). **(B)** Experimental binding curves showing FRET ratio measurements CaM proteins. For the experiments, FR was incubated either alone, or with a 30-fold excess of CaM^WT^, CaM^21A,57A^, CaM^94A,130A^, CaM^21A,57A,94A,130A^, in the presence of indicated [Ca^2+^] in 96-well plates. Emission at 480 nm and 535 nm was measured after excitation at 430 nm using a plate reader. The [Ex 430, Em 480] fluorescence intensity was then divided by the [Ex 430, Em 535] intensity to yield an emission ratio, which is plotted. Shown are mean ± standard deviation of 17 replicates for FR only, 15 replicates for CaM^WT^, 15 replicates for CaM^21A,57A^, 8 replicates for CaM^94A,130A^, and 3 replicates for CaM^21A,57A,94A,130A^.

To test these predictions experimentally, we used a plate reader to detect changes in FR-CaM binding as a function of free [Ca^2+^] with a 30-fold molar excess CaM^WT^, CaM^21A,57A^, CaM^94A,130A^, or CaM^21A,57A,94A,130A^ relative to FR. [CaM] was 673 nM (13 ng/μl), and [FR] was 22.9 nM (1.3 ng/μl). After collecting Ex 430/Em 480 and Ex 430/Em 535 data from different FR-CaM pairs (**Fig. S2**), we divided each [Ex 430/Em 480] fluorescence intensity value by its corresponding [Ex 430/Em 535] value to yield a ratio, which is plotted in **Fig. 3B**. We confirmed the presence of excess CaM compared to FR, and equal protein amount across experiments, by SDS-PAGE and SYPRO™ Ruby staining of the samples after FRET measurement (**Fig. S3**).

For the first model discrimination test, we compared the FRET ratio of CaM^WT^-FR pair with baseline in zero and low [Ca^2+^]. Contrary to the prediction of Model 1, we find no evidence that the signal in zero [Ca^2+^], or the area under the binding curve for 0 μM ≤ [Ca^2+^] ≤ 0.038 μM, is higher than baseline (*p* = 0.994 and *p* = 0.995, respectively, by a one-tailed Mann-Whitney *U* test). For the second test, we compared the FRET ratio of CaM^21A,57A^ -FR pair in high [Ca^2+^]. We found that FRET ratio in the maximum concentration used (39 μM [Ca^2+^]) is significantly higher than baseline (*p* = 2.83*10^−6^ by a one-tailed Mann-Whitney *U* test), as is the area under the complete binding curve (*p* = 2.83*10^−6^ by a one-tailed Mann-Whitney *U* test). These observations are sufficient to falsify Model 3, which predicts zero binding of MLCK to any CaM mutant with impaired N-terminus Ca^2+^ binding. As such, only the predictions of Model 2 are consistent with the full set of experimental results. As an additional test of Model 2, we examined binding of C-terminus mutant CaM^94A,130A^ in high [Ca^2+^]; consistent with the predictions of the model, we found no significant difference over baseline for the FRET ratio in 39 μM [Ca^2+^] or for the area under the complete binding curve (*p* = 0.345 and *p* = 0.294, respectively, by a one-tailed Mann-Whitney *U* test).

A striking feature of the experimental binding curves is the significant difference in maximum binding between CaM^WT^ and CaM^21A,57A^ (*p* = 1.16*10^−6^ by a one-tailed Mann-Whitney *U* test); this difference is also consistent with the predictions of Model 2 (**Fig. 3A**). Assuming Model 2, the fraction of CaM^21A,57A^ bound in the high-[Ca^2+^] limit is given by Eq. 2. For CaM^WT^, the limiting expression is the same as Eq. 2 but with K_2_ instead of K_4_ (for the reasons explained above), so that the model predicts equal maximum binding if K_2_ = K_4_. In the reference parameter set, K_2_ = 1,000 and K_4_ = 16.7, corresponding to predicted binding of 99.9% for CaM^WT^ and 92.0% for CaM^21A,57A^ in 39 μM [Ca^2+^]. As such, the experimentally observed difference is also predicted by our mathematical analysis, assuming that the literature parameter values are correct within an order of magnitude.

The validity of our experimental approach for model discrimination depends on the ability of the FR-CaM interaction to accurately mirror MLCK-CaM binding with sufficient sensitivity, as well as other considerations related to the experimental perturbations. To address these, we performed a series of control experiments and tested the robustness of our experimental design and data.

First, to confirm that the FRET ratio of FR is only altered by the binding of Ca^2+^-bound CaM, we collected 460 nm - 700 mm emission spectra with Ex 430 of the FR in 0 and 39 μM [Ca^2+^]. When FR was assayed alone, FR showed no changes in FRET ratio as a function of [Ca^2+^] and produced fluorescence peaks at Em 480 and Em 535 (**Fig. S4A**). Incubating FR with either CaM^WT^ or N-terminus mutant CaM^21A,57A^ in the presence of 39 μM Ca^2+^ caused FR fluorescence to increase at ∼Em 480 and decrease at ∼Em 535 (**Fig. S4B** and **S4C**). No change to FR fluorescence was observed in the absence of Ca^2+^ at any wavelength following FR incubation with CaM^WT^ or N-terminus mutant CaM^21A,57A^. C-terminus mutant CaM^94A,130A^ did not noticeably change the FR spectra at any wavelength, with or without Ca^2+^ (**Fig. S4D**).

Having observed no interactions between FR and CaM^94A,130A^ or CaM^21A,57A,94A,130A^ during our FRET-based binding assays, we wanted to confirm that the interactions we observed between FR-CaM^WT^ and FR-CaM^21A,57A^ are specific, Ca^2+^-dependent, and representative of binding between CaM and MLCK. To do so, we performed several on-bead binding assays using either FR or MLCK-FLAG. When bead-bound FR was incubated in 50 μl of buffer with 505 nM CaM^WT^ (approximately 20 ng/μl), the fraction of CaM^WT^ bound to FR increased proportionally with free [Ca^2+^]. At 39 μM [Ca^2+^], the majority of the CaM^WT^ input was bound to the FR (**Fig. 4A**). When the same experiment was repeated using N-terminus mutant CaM^21A,57A^, more than half of CaM^21A,57A^ bound to FR at 39 μM Ca^2+^ (**Fig. 4B**). These results suggest that the purified CaM proteins are functional and bind to Ca^2+^ and the FR, and that there is negligible non-specific binding between the FR and the purified CaM proteins. To determine if FR-CaM binding is representative of binding between CaM and full-length MLCK, we performed an on-bead binding assay after transient expression and purification of MLCK-FLAG protein. Despite the presence of several background bands, we observed the same CaM binding pattern to MLCK-FLAG, as to FR. At 39 μM [Ca^2+^], the majority of the CaM^WT^ input appeared bound to MLCK-FLAG (**Fig. S5A**), and there was partial binding of N-terminus mutant CaM^21A,57A^ to MLCK-FLAG (**Fig. S5B**). There was no apparent interaction between C-terminus mutant CaM^94A,130A^ and MLCK-FLAG at 39 μM [Ca^2+^] (**Fig. S5C**). None of the CaM proteins bound to MLCK-FLAG at 0 μM [Ca^2+^].

**Figure 4:**
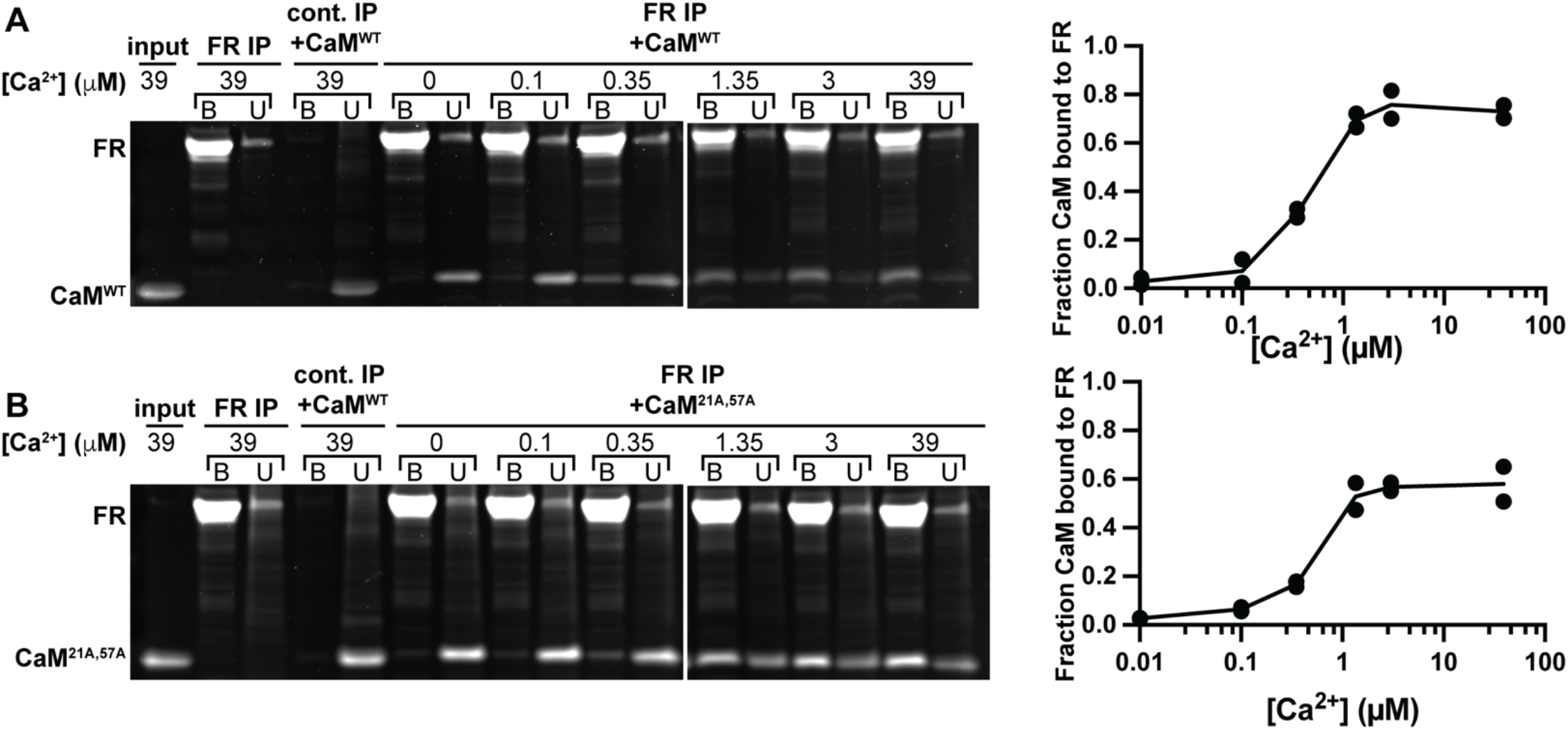
Purified CaM^WT^ and CaM^21A,57A^ show Ca^2+^-dependent interaction with FR. Representative images showing binding between bead-immobilized FR and **(A)** CaM^WT^ or **(B)** CaM^21A,57A^ in buffers with the indicated free [Ca^2+^]. Unbound and bound fractions were analyzed by SDS-PAGE followed by SYPRO™ Ruby protein gel stain. Fraction of CaM bound to the FR was quantified using ImageJ. The binding graph shows quantification of two experimental replicates; curve shows the average of the two data points at each [Ca^2+^]. FR without CaM addition and a control binding experiment using non-transfected HEK 293T cells serve as controls. The input lane shows the purified CaM^WT^ or CaM^21A,57A^ added to the on-bead binding assay.

We also investigated the relationship between the “fraction bound” calculated in the modeling analysis and the FRET ratio we use as a proxy for FR-CaM interaction. Although the Em 480/Em 535 ratio may not have a one-to-one relationship to “fraction bound” described in the models, this ratio correlates closely with FR-CaM binding at high and low [Ca^2+^], as calculated using on-bead binding assays **(Fig. S6)**. Moreover, the same FRET ratio has been shown to mirror MLCK phosphorylation *in vivo* (*14*). As a result, we conclude that the ratio can be used to characterize CaM-FR interaction for our model discrimination strategy, which rests on differential binding predictions in zero and high [Ca^2+^].

We do not observe binding between FR and C-terminus mutant CaM^94A,130A^ in our FRET-based binding assays. To investigate the possibility that CaM^94A,130A^ might bind to the FR but fail to interfere with FRET, we used a Bio-Layer Interferometry (BLI) assay as an orthogonal method to measure binding between FR and CaM. Binding detection in the BLI assay is independent of FRET measurement, which enabled us to decouple binding and FRET interference. The BLI assay showed robust binding between FR and CaM^WT^ at high [Ca^2+^], an intermediate degree of binding between FR and CaM^21A,57A^ at high [Ca^2+^], and no detectable binding between FR and CaM^94A,130A^ at high [Ca^2+^] (**Fig. 5A**). Performing a BLI assay using CaM^WT^, CaM^21A,57A^ and CaM^94A,130A^ with purified MLCK-FLAG yielded comparable results (**Fig. 5B**).

**Figure 5:**
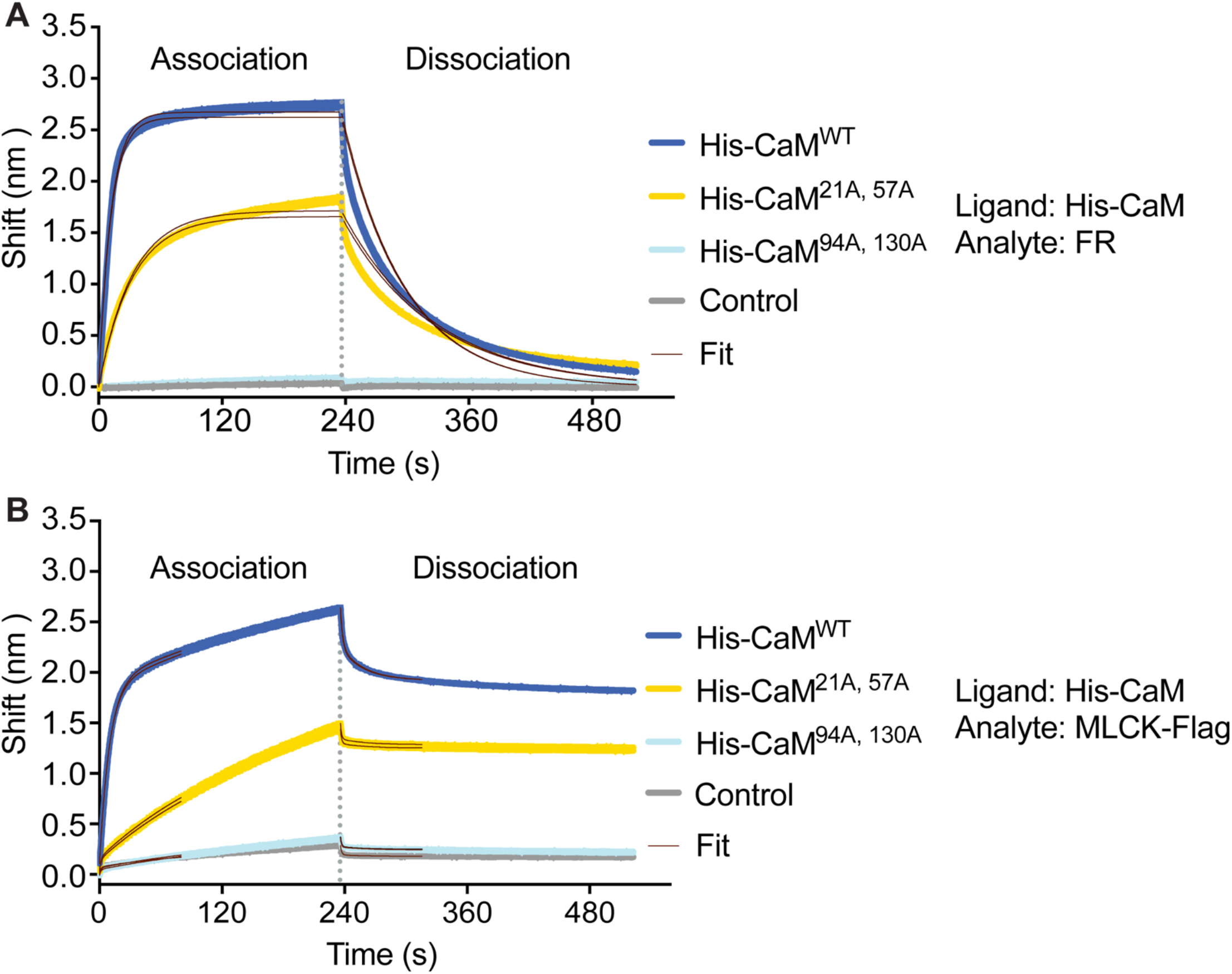
Bio-Layer Interferometry (BLI) binding assays show no binding between FR or MLCK-FLAG and CaM^94A,130A^. BLI was performed using purified His-tagged calmodulin (CaM^WT^, CaM^21A,57A^, or CaM^94A,130A^) as a ligand immobilized to a Ni-NTA biosensor, and either **(A)** FR or **(B)** MLCK-FLAG as an analyte. Two replicates of each condition are shown. “Control” refers to the analyte binding to the ligand-free probe.

We next assessed the intrinsic sensitivity of the FRET assay. To test if our assay can detect small changes in binding, which is necessary for many of the comparisons involved in our model discrimination analysis, we measured FRET at 39 μM [Ca^2+^] with different FR:CaM ratios. Assuming Model 2 and the reference parameter values from Fajmut, Brumen *et al*. (*9*), if the FR:CaM molar ratio is increased from 2:1 to 2:1.2, the amount of CaM-bound FR in 39 μM [Ca^2+^] is predicted to increase by 8.5% (from 46.2% bound to 54.7% bound). When we repeated our FRET-based FR-CaM binding assay using these ratios (22.9 nM FR and 11.45 nM or 13.75 nM CaM), we observed a significant increase in F480/F535 **(Fig. S7)** (*p* = 0.00029 by a one-tailed Mann-Whitney *U* test), with a calculated 8.7% difference in FRET ratios between the two conditions. We therefore conclude that, under our experimental conditions, the assay is sensitive enough to detect at least a 9% increase in the fraction of CaM-bound FR.

The predictions of zero MLCK binding in a particular [Ca^2+^] hold for any set of numerical values assigned to the model parameters (the equilibrium constants K_1_…K_11_ in **Fig. 1)**, as shown in Dexter *et al*. (*12*). For predictions of non-zero binding, however, the magnitude of binding does depend on the choice of parameters. As such, it is theoretically possible that the model could predict a binding fraction that is non-zero but too small to detect experimentally in an important concentration regime. To address this potential concern, we undertook a sensitivity analysis of the two key predictions of non-zero MLCK binding. For numerical calculations with the models, we used the set of reference parameter values compiled by Fajmut, Brumen *et al*. from previous biochemical studies (*9*). As shown in Eq. 1, the prediction of Model 1 for the fraction of MLCK bound in zero [Ca^2+^] depends on the value of a single parameter (K_9_). Assuming the parameter value selected by Fajmut, Brumen *et al*. (*9*) based on several prior studies (K_9_ = 0.078 μM^-1^) and the concentrations of CaM^WT^ and FR used in the main experiment, the model predicts 5.2% binding in zero [Ca^2+^]. As is clear from the structure of Eq. 1, the fraction of MLCK predicted to bind in zero [Ca^2+^] increases with the ratio of total CaM to total MLCK. To confirm our falsification of Model 1, we therefore repeated the binding experiment in zero [Ca^2+^] with a much higher [CaM] (35.65 μM), for which 73.5% binding is predicted with the reference parameter values and 21.7% binding is predicted with K_9_ = 0.0078 μM^-1^ (i.e., 10-fold lower than the reference value). As in the main experiment, the FRET ratio did not increase above baseline in zero [Ca^2+^] (**Fig. S8**), providing strong evidence against Model 1 even if previous parameter estimates are incorrect by an order of magnitude (*p* =0.998 by a one-tailed Mann-Whitney *U* test).

For Model 2, the fraction of MLCK predicted to bind to the N-terminus mutant CaM^21A,57A^ depends on two parameters, K_4_ and K_6_ (Eq. 2). We confirmed the robustness of the key prediction of non-zero binding in 39 μM [Ca^2+^] by calculating 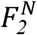 for 200,000 combinations in which values for each of the two parameters were chosen at random from the interval [0.1v, 10v], where v is the reference value from Fajmut, Brumen *et al*. (*9*). Across the combinations 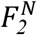 was never less than 18%, a level of binding straightforward to distinguish from baseline; the full distribution is shown in **Fig. S9**.

Finally, we confirmed that the addition of the purified proteins to the prepared Ca^2+^ buffers did not significantly alter the buffers’ free [Ca^2+^]. We measured the fluorescence intensity of two Ca^2+^ indicator dyes with different Ca^2+^ affinities (Fluo-4 and Calcium Green), before and after addition of the buffer in which our CaM proteins were stored. The largest volume of protein added to any of our FRET-based binding experiments was 1.3 μl per assay, so this was the volume that we tested. We did not observe a significant change in indicator dye fluorescence intensity after CaM storage buffer addition, which suggests that any changes to free [Ca^2+^] that are experimentally introduced are smaller than can be detected reliably with Fluo-4, a dye that has a very high Ca^2+^ affinity (17 nM) **(Fig. S10)** (*p* =0.23-0.99 by a two-tailed Mann-Whitney *U* test).

These sets of control experiments and sensitivity analyses show that our experimental system is robust and rigorously controlled. The FRET-based binding assay that we used as the core of our model discrimination strategy is well validated. Furthermore, the FR and CaM proteins we used are free of major contaminants. SYPRO® stained SDS-PAGE gels of the CaM proteins also confirm that these proteins display the predicted altered electrophoretic mobility upon binding to Ca^2+^, which strongly suggests that they bind [Ca^2+^] as expected. Follow-up plate reader experiments confirmed that FRET interference is sensitive enough to allow discrimination between candidate CaM-MLCK binding models. We also show that a change in FRET ratio is caused only by binding of Ca^2+^-bound CaM proteins. Conversely, orthogonal BLI experiments confirm that a lack of FRET interference accurately reflects a lack of FR-CaM binding. We have also confirmed that the [Ca^2+^] of our experimental buffers is not altered by the addition of our experimental proteins. Critically, on-bead binding experiments with both FR and MLCK-FLAG reproduced the key findings of our FRET-based binding assays. These data, in combination with the MLCK-FLAG data from our BLI assays, suggests that FR-CaM binding accurately mirrors binding between CaM and full-length MLCK protein, and suggests that our FR data can confidently be used for our model discrimination strategy.

In sum, our binding measurements using the FR falsify key predictions of both the fully random binding mechanism (Model 1) and the fully ordered binding mechanism (Model 3). Our data instead support Model 2, which predicts partially ordered binding between CaM and MLCK (**Fig. 1**). As predicted by Model 2, we observe zero binding in low [Ca^2+^] for CaM^WT^ and all CaM mutants, non-zero binding in high [Ca^2+^] for CaM^WT^, non-zero binding in high [Ca^2+^] for N-terminus mutant CaM^21A,57A^, and zero binding in high [Ca^2+^] for both of our CaM mutants containing mutations in the C-terminus domain (CaM^94A,130A^ and CaM^21A,57A,94A,130A^). Model 2 makes six correct predictions and no incorrect predictions (**Table 1**). Model 1 makes two falsified predictions: non-zero binding in low [Ca^2+^] for CaM^WT^, and non-zero binding in high [Ca^2+^] for the C-terminus mutant, CaM^94A,130A^. Finally, Model 3 makes one falsified prediction: zero binding in high [Ca^2+^] for the N-terminus mutant, CaM^21A,57A^.

**Table 1:** Summary of model discrimination results for CaM and MLCK binding. B denotes binding, NB denotes no binding. Bolded entries indicate experimentally falsified predictions.

|  | model 1 | model 2 | model 3 | observed |
| --- | --- | --- | --- | --- |
| *zero Ca<sup>2+</sup>* |  |  |  |  |
| CaM<sup>WT</sup> | **B** | NB | NB | NB |
| CaM<sup>21,57A</sup> | **B** | NB | NB | NB |
| CaM<sup>94A,103A</sup> | **B** | NB | NB | NB |
| *high Ca<sup>2+</sup>* |  |  |  |  |
| CaM<sup>WT</sup> | B | B | B | B |
| CaM<sup>21,57A</sup> | B | B | **NB** | B |
| CaM<sup>94A,103A</sup> | **B** | NB | NB | NB |

## Discussion

In recent work, Dexter *et al*. analyzed a class of previously developed theoretical models of CaM-MLCK binding and proposed a multi-part strategy for distinguishing between them (*12*). Our primary contribution here is an experimental implementation of this model discrimination strategy, which suggests a partially ordered mechanism for binding. Our analysis sheds new light on a controversy that has persisted for several decades and demonstrates a productive interplay between mathematical modeling and biochemical analysis.

It is important to remember that model discrimination strategies of this kind work by process of elimination. Models can be ruled out when their predictions contradict experimental data, but the failure of some models does not guarantee the correctness of others. We present here evidence sufficient to falsify all but one of the models of CaM-MLCK binding in the literature (**Table 1**). Data collected using wild-type and mutant CaM proteins in three different binding experiments (FRET, BLI and on-bead binding) in zero and high [Ca^2+^], using both FR and MLCK-FLAG when possible, rule out both Model 1 and Model 3. Our data is consistent with Model 2, which makes six correct predictions. Taken together, these results strongly support a partially ordered mechanism in which MLCK can bind to CaM after Ca^2+^ is bound at the C-terminus (Model 2).

To investigate MCLK-CaM binding, we centered our experimental design on a FR system that uses the CaM-binding domain of smooth muscle MLCK. Prior studies have demonstrated that this reporter accurately reflects the MLCK-CaM binding in a variety of *in vivo* and *in vitro* contexts (*13-19*). Moreover, our FRET-based FR-CaM binding assay enabled collection of robust and reproducible data using purified CaM and FR proteins, with minimal risk of interference from endogenous proteins or contaminants. This system is also amenable to complementary, non-FRET-based biochemical techniques to assess FR-CaM binding, which allowed us to confirm the FRET-based binding data we obtained. Although the FR we used has been extensively validated to be a good proxy for MLCK-CaM binding, the use of an MLCK fragment still does raise concerns about the behavior of the fragment relative to the full-length protein. To allay these concerns, we repeated two key experiments – on-bead binding experiments and BLI – with a full-length MLCK-FLAG protein in lieu of the FR protein. In each of these experiments, FR and MLCK-FLAG performed similarly, showing the utility of the FRET-based binding assay for our model discrimination strategy.

In HEK 293T lysates, changing myosin phosphorylation has been shown to correspond to changes in the FRET ratio produced by FR. Changes to the FRET ratio can therefore be used as a proxy for MLCK activation (*14*). It is important to note that changes in FRET ratio do not necessarily correspond 1:1 with changes to fraction of FR bound to CaM across all of the Ca^2+^ concentrations. Nevertheless, this fact does not impede our model discrimination strategy. The FRET ratios measured at 0 μM [Ca^2+^] and 39 μM [Ca^2+^] are the data points most critical to our model discrimination strategy. We also complement this FRET-based binding assay data with on-bead binding assays showing that in 39 μM [Ca^2+^], the majority of CaM^WT^ binds to either FR or full length MLCK protein (MLCK-FLAG) expressed using human smooth muscle MLCK gene *MYLK1*, while a negligible amount of CaM binds to FR or MLCK-FLAG in 0 μM [Ca^2+^]. We therefore conclude that FRET ratio data represents MLCK-CaM^WT^ binding in 0 μM [Ca^2+^] and 39 μM [Ca^2+^].

It should be noted that mammalian myosin light chain kinases are a group of serine/threonine kinases encoded by at least four genes: *MYLK1, MYLK2, MYLK3*, and *MYLK4* (*26*). *MYLK1* has alternative initiation sites that enable expression of at least four protein products including nonmuscle (long isoform) MLCK and smooth muscle (short isoform) MLCK. *MYLK2* encodes an MLCK isoform expressed solely in skeletal muscle, *MYLK3* encodes a cardiac-specific MLCK (MLCK3), and the gene product(s) of *MYLK4* remain largely uncharacterized (*25, 26*). All MLCK proteins, except MLCK4, have a CaM-binding peptide that shows sequence homology to both the peptide used in our FR, which is derived from avian smooth muscle MLCK, and to the CaM-binding domain of our MLCK-FLAG protein. As a result, we expect that our findings here can be generalized to other MLCK proteins and their interaction with CaM and Ca^2+^.

In addition to providing a blueprint for future model discrimination efforts that integrate mathematical and biochemical approaches, our work may also prove useful in a translational or pharmaceutical context, as MLCK and its activation by CaM have been linked to the pathogenesis of human disease. For example, increased organization of the sarcomere, the contractile unit of the striated muscle, is observed during the onset of cardiomyocyte hypertrophy. CaM-activated MLCK has been shown to mediate sarcomere organization induced by a hypertrophic agonist in cultured cardiomyocytes and *in vivo* (*27*). Fuller characterization of the Ca^2+^-CaM-MLCK interaction network may therefore prove relevant for future drug discovery efforts.

## METHODS and MATERIALS

### Resource availability

Further information and requests for reagents and resources may be directed to and will be fulfilled by corresponding author Dr. Yasemin Sancak.

### Experimental model and subject details

HEK 293T cells were acquired from the Sabatini Lab at the Whitehead Institute for Biomedical Research. The identity of the cell line was confirmed using STR analysis. Cells were grown in DMEM supplemented with 1X GlutaMAX and 10% fetal bovine serum. Cells were tested for mycoplasma every 3 months and were confirmed to be free of mycoplasma contamination. Cells were cultured at 37°C with 5% CO2.

#### Cell Culture

- DMEM; Thermo Fisher Scientific, cat. no. 11-965-118
- Glutamax; Fisher Scientific, cat. no. 35-050-061
- PBS; Thermo Fisher Scientific, cat. no. 20012050
- Penicillin/streptomycin solution; VWR cat. no. 45000-652
- Fetal bovine serum; Life Technologies, cat. no. 26140087
- Genlantis MycoScope PCR Detection Kit; VWR cat. no. 10497-508

#### Gel Sample Preparation, Running, and Imaging

##### Reducing sample buffer, pH 6.8

- SDS; Sigma-Aldrich Cat. no. L4509-1KG
- BME/2-mercaptoethanol; Sigma-Aldrich Cat. no. M3148-25ML
- Glycerol; Sigma-Aldrich Cat. no. G5516-1L
- Tris–HCl: Sigma-Aldrich cat. no. T5941
- Bromophenol Blue; VWR Cat. no. 97061-690

##### Gel Electrophoresis

- 10X Tris/Glycine Buffer; Boston BioProducts Cat. no. BP-150-4L
- Tris-Glycine 12% Gel, 10-well or 15-well; Bio-Rad Cat. No. 4561043 or 4561046

##### Gel Imaging

- Methanol; Sigma-Aldrich cat. no. 32213-2.5L
- Acetic Acid; Sigma-Aldrich cat. no. A6283-500ML

Gels were stained with SYPRO® Ruby Protein Gel Stain (# S12000) following manufacturer’s instructions and imaged with iBrightCL 1000 on fluorescent protein gel setting.

#### Expression of FR and MLCK-FLAG via transient transfection

- Transfection reagent; X-treme(GENE) 9, Sigma-Aldrich, cat. no. 6365779001

Day 1: Either 5 million (purification) or 2 million (on-bead binding assays) HEK 293T cells were plated on 15 cm or 10 cm plates, respectively. Day 2: 5 μg FR plasmid (purification), 1.5 μg FR plasmid (on-bead binding assays with FR) or 3 μg MLCK-FLAG plasmid (on-bead binding assays with MLCK-FLAG) was transfected using transfection reagent. *Day 4*: Cells were harvested, and FR or MLCK-FLAG was either purified (see below, “Purification of FR”) or used for on-bead binding assays.

#### Purification of FR and MLCK-FLAG

##### Lysis Buffer

- 50 mM HEPES-KOH (HEPES, Sigma-Aldrich, # H3375-1KG KOH, Sigma Millipore, # 1050121000)
- 150 mM NaCl, Sigma-Aldrich, # 746398-5KG
- 5 mM Ethylenediaminetetraacetic acid (EDTA): Sigma-Aldrich, # 607-429-00-8
- 1% Triton X-100: Sigma, # X100-1L
- FLAG Peptide Elution Buffer: 50 mM HEPES, 500 mM NaCl, pH 7.4
- Protease Inhibitor Tablets; Complete™, Mini, EDTA-free Protease Inhibitor Cocktail, Sigma-Aldrich cat. no. 5892953001
- Anti-FLAG affinity gel; Sigma-Aldrich, cat. no. A2220-5ML
- 3X FLAG peptide; Sigma-Aldrich cat. no. F4799-4MG
- Chromatography spin column; Bio-Rad cat. no. 7326204

Cells were lysed using lysis buffer supplemented with proteases inhibitors. Lysates were triturated in tubes and then centrifuged at 17000 g for 10 minutes. Cell supernatant was divided into three tubes with 200 μl of anti-FLAG affinity gel slurry (50:50 bead/lysis buffer). These tubes were rocked at 4 °C for 1 hour, and beads were washed three times with lysis buffer. A 22.5-gauge syringe was used to aspirate all remaining liquid from the tubes. FLAG-tagged protein was eluted with 90 μl of elution buffer, and 10 μl of 3XFLAG peptide prepared at 5 mg/mL for 30 minutes at 30 °C. The gel/elution buffer slurry from all three tubes was then pipetted into one spin column and spun at max speed for 5 minutes.

#### Expression and Purification of Wild-type and Mutant CaM via Bacterial Induction

- Low-salt LB broth: Millipore Sigma, # L3397-1KG
- Isopropyl beta-D-1-thiogalactopyranoside (IPTG): Sigma-Aldrich, # I5502-5G
- HiTrap Q HP anion exchange chromatography column: Cytiva, # 17115301
- Centrifugal protein concentrator, 10K cutoff: Amicon, # UFC8010

##### Lysis Buffer

- 20 mM Tris-HCl, Sigma-Aldrich cat. no. T5941
- 100 mM NaCl, Sigma-Aldrich, cat. no. 746398-5KG
- 20 mM imidazole, Sigma-Aldrich, cat. no. I5513
- 0.4 mM EGTA, Sigma-Aldrich cat. no. 324626-25GM
- 0.5 mM TCEP, Sigma-Aldrich cat. no. C4706-2G

##### Elution Buffer

- 20 mM Tris-HCl, Sigma-Aldrich cat. no. T5941
- 100 mM NaCl, Sigma-Aldrich, cat. no. 746398-5KG
- 200 mM imidazole, Sigma-Aldrich, cat. no. I5513
- 0.4 mM EGTA, Sigma-Aldrich cat. no. 324626-25GM
- 0.5 mM TCEP, Sigma-Aldrich cat. no. C4706-2G

##### Buffer A

- 100 mM NaCl, Sigma-Aldrich, cat. no. 746398-5KG
- 20 mM Tris-HCl, Sigma-Aldrich cat. no. T5941

##### Buffer B

- 1M NaCl, Sigma-Aldrich, cat. no. 746398-5KG
- 20 mM Tris-HCl, Sigma-Aldrich cat. no. T5941

BL21 (DE3) bacteria transformed with a CaM expression vector were grown at 37 °C, 200 RPM until OD600 reached 0.8-1.0. Cultures were cooled to 18 °C and supplemented with IPTG to a final concentration of 0.3 mM before incubation at 18 °C, 200 RPM for 18 hours. Bacteria were pelleted, frozen, and stored at -80 °C until protein purification. Cells were lysed using sonication for 5 minutes at 40% power before being centrifuged at 17,400 RPM for 40 minutes to separate the soluble and insoluble portions of the lysate. The clarified lysate was loaded onto a Ni-NTA gravity column and then washed with 10 column volumes of lysis buffer before elution with 5 column volumes of elution buffer. The eluted protein was loaded manually into a 1 ml anion exchange chromatography column. HPLC was performed using buffers A and B. The HPLC fractions that contained the highest concentrations of protein were collected and concentrated to their final concentrations using a centrifugal filter with a 10 kDa molecular weight cutoff. Stock protein concentrations were as follows: CaM^WT^: 3.98 μg/μl; CaM^21A,57A^: 4.96 μg/μl; CaM^94A,130A^: 1.36 μg/μl; CaM^21A,57A,94A,130A^: 1.52 μg/μl. Purified protein was aliquoted, flash frozen in a liquid nitrogen dewar, and then stored at -80 °C until use.

#### FRET-Based CaM-FR Binding Assay

- Calcium calibration buffer kit; Invitrogen cat. no. C3008MP
- Microplate reader; BioTek Synergy H1
- Black 96-well plates; Greiner Bio-One cat. no. 655076

For each CaM-FR binding assay condition, 2 μg of CaM (either wild-type or mutant) and 200 ng FR was thoroughly mixed with 150 μl of Ca^2+^ buffer (14 conditions between 0 μM and 39 μM). 150 μl of the resulting mixture was pipetted into 1 well of a 96-well plate (Greiner Bio-One, # 655076). CaM was used at a final concentration of 673 nM (13 ng/μl); FR was used at a final concentration of 22.9 nM (1.3 ng/μl). All protein preps (FR and CaM) were at sufficiently high concentrations that each protein addition added negligible volume; stock protein concentrations were as follows: CaM^WT^: 3.98 μg/μl; CaM^21A,57A^: 4.96 μg/μl; CaM^94A,130A^: 1.36 μg/μl; CaM^21A,57A,94A,130A^: 1.52 μg/μl; FR: 0.52 μg/μl. Follow-up experiments with Ca^2+^ indicator dyes confirmed that the final [Ca^2+^] were unaffected by the addition of CaM or FR to the Ca^2+^ buffer (see “CaM and Ca^2+^ Buffer with Ca^2+^ Indicator Dye Assay” below). For functional assay, all 14 Ca^2+^ conditions were tested together, batch-wise. All wells were read at 430, 480 excitation/emission, and then again at 430, 535 excitation/emission using a microplate reader. The [430, 480] read was divided by the [430, 535] read to yield an emission ratio, which was plotted relative to [Ca^2+^]. A small portion of each prepared Ca^2+^ condition was conserved and prepared with sample buffer, to be loaded onto a gel and stained.

#### CaM-FR FRET-Based Binding Sensitivity Assays

- Microcon concentrator column, Millipore cat. No. 42407

To test the robustness of the falsification of Model 1 to uncertainty in parameter values, we repeated the binding assay in zero [Ca^2+^] using 35.65 μM CaM^WT^. CaM^WT^ was concentrated to a stock concentration of 66.6 μg/μl using a concentrator column so that a comparable volume of protein stock (relative to previous assays) could be used. The “FRET-Based CaM-FR Binding Assay” was then performed as described above, except using 35.65 μM CaM^WT^.

To determine a lower sensitivity limit for the CaM-FR binding assay, 22.9 nM FR was added to 150 μl of high Ca^2+^ buffer along with either 11.45 or 13.74 nM CaM^WT^, which resulted in a CaM^WT^:FR molar ratio of either 1:2 or 1.2:2. 150 μl of the resulting mixture was then read in a plate reader using the program employed by the “FRET-Based CaM-FR Binding Assay” (see above) and analyzed accordingly.

#### CaM-FR Spectral Scanning

- Calcium calibration buffer kit; Invitrogen cat. no. C3008MP
- Microplate reader; BioTek Synergy H1
- Black 96-well plates; Greiner Bio-One cat. no. 655076

CaM and FR were added to either 0 or 39 μM [Ca^2+^] as per “FRET-Based CaM-FR Binding Assay” above. CaM was used at a final concentration of 673 nM (13 ng/μl); FR was used at a final concentration of 22.9 nM (1.3 ng/μl). Using an excitation wavelength of 430 nm, the wells were read using spectral scanning between wavelengths 460 nm and 700 nm, with an emission step of 10 nm. The fluorescence intensity of each wavelength was then plotted.

#### CaM-FR and CaM-MLCK-FLAG On-Bead Binding Assays

For assays utilizing FR, assay data was collected under 8 different conditions: FR bound to anti-FLAG M2 affinity gel beads and CaM under 6 different [Ca^2+^] (0 μM, 0.1 μM, 0.35 μM, 1.35 μM, 3 μM, and 39 μM) and two controls (bead-bound FR alone in 39 μM [Ca^2+^]; beads with CaM in 39 μM [Ca^2+^]). For assays utilizing MLCK-FLAG, assay data was collected under 4 different conditions: MLCK-FLAG bound to anti-FLAG M2 affinity gel beads and CaM under 2 different [Ca^2+^] (0 μM and 39 μM) and two controls (bead-bound MLCK-FLAG alone in 39 μM [Ca^2+^]; beads with CaM in 39 μM [Ca^2+^]).

##### Binding Assay Protocol

Transiently transfected cells were washed once with chilled PBS, and then harvested in 1% Triton buffer using a cell scraper. Cells were triturated to ensure complete lysis, and then were spun down at maximum speed (17,000g) for 10 minutes using a 4 °C centrifuge. 160 μl bead:glycerol M2 FLAG slurry was prepared for FLAG-tagged protein binding by washing three times with 1 ml 1% Triton buffer before being divided evenly between 8 tubes. The cleared lysate from the transiently transfected cells was divided evenly between 7 of the tubes, while cleared lysate from control WT HEK 293T was added to the 8^th^ tube (control lysate). The beads were incubated with the lysate on a rocker at 4 °C for 45 minutes. After incubation, the two control tubes were washed twice with 200 μl 39 μM [Ca^2+^], while the tubes for the 6 different Ca^2+^ conditions were each washed with the appropriate Ca^2+^ buffer. 50 μl of the appropriate Ca^2+^ buffer was then added to each experimental tube, while 50 μl of 39 μM Ca^2+^ was added to each control tube. Finally, for FR assays, 0.5 μg of CaM was added to all tubes except the reporter-only condition; final CaM concentration was approximately 505 nM (10 ng/μl). For MLCK-FLAG assays, 0.25 μg of CaM was added to all tubes except the MLCK-FLAG-only condition; final CaM concentration was approximately 24 nM (5 ng/μl) (NOTE: CaM concentration for MLCK-FLAG assays was decreased to compensate for decreased MLCK-FLAG expression relative to FR expression). CaM stock was at a sufficiently high concentration to ensure the addition of CaM did not meaningfully alter Ca^2+^ buffer concentration (see “CaM and Ca^2+^ Buffer with Ca^2+^ Indicator Dye Assay below”) (**Fig. S5**). All tubes were incubated at room temperature for 15 minutes and were flicked occasionally. Following incubation, the tubes were spun for 1 minute at 1,000 g to pellet the beads. 40 μl of the unbound portion was conserved and prepared with 10 μl 5X reducing sample buffer. The beads for all conditions were then washed with 50 μl of the appropriate Ca^2+^ buffer and all liquid was aspirated from the beads using a syringe to remove residual unbound CaM. The beads were then incubated with 40 μl of FLAG peptide prepared in elution buffer for 30 minutes at 30°C to elute FLAG tagged FR. After elution, the beads were loaded to a spin column and spun for 1 minute at 1000g at room temperature to recover the eluate without the beads. This eluted sample was then boiled with 10 μl 5X reducing sample buffer. Finally, an “input” sample was prepared by adding 0.5 μg of CaM to 40 μl of 39 μM [Ca^2+^] solution and 10 μl 5X reducing sample buffer.

##### Binding quantification

Bands were quantified using ImageJ. A rectangle was drawn around the unbound CaM band in the FR / CaM assay at 0 μM [Ca^2+^]. This same rectangle was used to quantify the mean gray value within the rectangle for each binding condition. A measurement was also taken of the gel background, from the “input” lane containing only calmodulin. The background intensity measurement was subtracted from each quantified region. % binding was calculated using the following equation: % binding = bound fraction / [unbound CaM fraction + bound CaM fraction]. For **Fig. S6**, binding fraction was normalized by setting binding at 0 μM [Ca^2+^] to 0% and at 39 μM [Ca^2+^] to 100%, and calculating the % bound in the rest of the samples by normalizing to this scale.

##### CaM and Ca^2+^ Buffer with Ca^2+^ Indicator Dye Assay

- Calcium Green 5N, Hexapotassium Salt, Cell Impermeant; Thermo Fisher Scientific, cat. no. C3737
- Fluo-4 Cell Impermeant; Thermo Fisher Scientific, cat. no. F14200
- Microplate reader; BioTek Synergy H1
- Calcium calibration buffer kit; Invitrogen cat. no. C3008MP
- Black 96-well plates; Greiner Bio-One cat. no. 655076

##### CaM Storage Buffer (“Dummy Buffer”)

- 250 mM NaCl, Sigma-Aldrich, cat. no. 746398-5KG
- 20 mM Tris, pH 8, Sigma-Aldrich cat. no. T5941

This assay was performed to confirm that the addition of CaM protein to the Ca^2+^ buffer used for the CaM-FR binding assay did not change the final [Ca^2+^]. Two “sets” of each of the 14 different Ca^2+^ buffers used for CaM-FR binding assay were prepared, with dye (Calcium Green or Fluo-4) added to a final concentration of 1 μM. For one set of buffers, the buffer in which the CaM proteins was stored was added (“dummy buffer”). The amount of “dummy buffer” added was equal in volume to the maximum volume of CaM added per reaction (ie the C-terminus mutant CaM was the most dilute CaM prep, and approximately 1.5 μl CaM was added to each assay well, so an equivalent amount of “dummy buffer” was used for these experiments). 150 μl of each of these prepared buffers was added to wells of a black 96-well plate and data was collected using the GFP filter of a plate reader.

#### Mass Spectrometry Analysis of CaM Proteins

- Calcium calibration buffer kit, Invitrogen cat. no. C3008MP
- Chloroacetamide, Sigma cat. no. C0267
- TCEP, Sigma-Aldrich cat. no. C4706-2G
- Rappsilbers Stage tipping Paper (23), C-18 material: CDS Empore C18 Extraction Disks, Fisher cat. no. 13-110-016
- Formic acid for HPLC LiChropur, Sigma cat. no. 5438040100
- Trifluoroacetic acid (TFA), HPLC Grade, 99.5+%, Alfa Aesar, cat. no. AA446305Y
- Acetonitrile (ACN), Sigma-Aldrich cat. no. 271004-100ML
- Mass spectrometer: LC-MS: Orbitrap Fusion Lumos Tribrid (Thermo Fisher Scientific) and EASY-nLC™ 1200 System (Thermo Fisher Scientific)
- Triethylammonium bicarbonate buffer 1.0 M, pH 8.5±0.1, Sigma cat. no. T7408
- Ethanol for HPLC, Sigma cat. no. 459828
- Promega trypsin, cat no. V5111
- Methanol for HPLC, Sigma cat. No. 494291

##### StageTip Buffer A

- 5% ACN
- 0.1% TFA
- MQ water

##### StageTip Buffer B

- 0.1% TFA
- 80% ACN
- MQ water

##### Extraction solution

- 0.1% TFA, diluted in MQ water

##### Running the Gel

30 μg of wild-type CaM and N-terminus mutant CaM were each mixed with high Ca^2+^ calibration buffer to a final volume of 20 μl. A blank control consisting of CaM storage buffer and Ca^2+^ calibration buffer was also prepared. All samples were reduced using 1 mM final concentration of TCEP and alkylated using 2 mM final concentration of chloroacetamide (CAM) at 37C for 20 min on a shaker. They were then quenched using 1 mM final concentration of TCEP at 37C for 20 min on a shaker. Next, they were mixed with 5X gel loading buffer, boiled for five minutes, and loaded into a 10% gel (see “**Gel Electrophoresis**” above). The samples were run until the dye front reached the bottom of the gel. The gel was then rinsed and stained using Coomassie stain for one hour, and destained overnight.

### In-gel Digestion Protocol

The next day, two bands from each lane were excised and cut into small cubes using a single-use scalpel; one band was comprised of the gel at ∼18 kDa (where the major CaM band is located) and one was comprised of the gel between ∼10 and 17 kDa, the area under the major band. The rest of the in-gel digestion was performed as previously described in (*28*).

### Extraction of Peptides and StageTip Purification

StageTip extraction of peptides was performed as described previously in (*29*).

### nanoLC-MS/MS Analyses

LC-MS analyses were performed as described previously with the following minor modifications (*30*). Peptide samples were separated on a EASY-nLC 1200 System (Thermo Fisher Scientific) using 20 cm long fused silica capillary columns (100 μm ID) packed with 3 μm 120 Å reversed phase C18 beads (Dr. Maisch, Ammerbuch, DE). The LC gradient was 90 min long with 5−35% B at 300 nL/min. LC solvent A was 0.1% (v/v) aq. acetic acid and LC solvent B was 20% 0.1% (v/v) acetic acid, 80% acetonitrile. MS data was collected with a Thermo Fisher Scientific Orbitrap Fusion Lumos Tribrid mass spectrometer. Data-dependent analysis was applied using Top15 selection with CID fragmentation

### Computation of MS raw files

Data .raw files were analyzed by MaxQuant/Andromeda (*31*) version 1.5.2.8 using protein, peptide, and site FDRs of 0.01 and a score minimum of 40 for modified peptides, 0 for unmodified peptides; delta score minimum of 17 for modified peptides, 0 for unmodified peptides. MS/MS spectra were searched against the UniProt human database (updated July 22nd, 2015). MaxQuant search parameters: Variable modifications included Oxidation (M) and Phospho (S/T/Y). Carbamidomethyl (C) was a fixed modification. Maximum missed cleavages was 2, enzyme was Trypsin/P and max. charge was 7. The MaxQuant “match between runs” feature was disabled. The initial search tolerance for FTMS scans was 20 ppm and 0.5 Da for ITMS MS/MS scans.

### Semi-quantitative analysis of contaminants and CaM protein

The intensity of peptides annotated as CaM, keratin, or bacterial protein was summed up and their relative intensities were calculated by dividing them to the total peptide intensity in the samples.

#### Octet BioLayer Interferometry (BLI) measurement

- Octet Red 96 (ForteBio, Pall Life Sciences)
- Bovine Serum Albumin (BSA), Sigma-Aldrich cat. no. A3294-100G
- Calcium chloride, Sigma-Aldrich cat. no. 746495-1KG
- Tris-HCl, Sigma-Aldrich cat. no. T5941
- NaCl Sigma-Aldrich, # 746398-5KG

The binding of His-CaM to FR or FL-MLCK-FLAG protein in the presence of Ca^2+^ was analyzed using the Octet Red 96 (ForteBio, Pall Life Sciences) following the manufacturer’s procedures in duplicates. The reaction was carried out in black 96 well plates maintained at 30 °C. The reaction volume was 200 μL in each well. The Octet buffer contained 20 mM Tris-HCl, 200 mM NaCl, and 0.1% BSA, pH 8.0. The Association buffer contained 20 mM Tris-HCl, 200 mM NaCl, 1.5 mM Ca^2+^ and 0.1% BSA, pH 8.0. The Dissociation buffer contained 20 mM Tris-HCl, 200 mM NaCl, and 0.1% BSA, pH 8.0. The concentration of ligand – His-CaM^WT^, His-CaM^21A, 57A^, or His-CaM^94A, 130A^ – in the Octet buffer was 2 μM. The concentration of His-GST as the quench in Octet Buffer was 0.68 μM. The concentration of FR or MLCK-FLAG as the analyte in the Association buffer was 1.5 μM. Ni-NTA optical probes were loaded with His-CaM^WT^, His-CaM^21A, 57A^, or His-CaM^94A, 130A^ as ligands for 110 seconds and quenched with His-GST for 80 seconds prior to binding analysis. While not loaded with ligand, the control probes were quenched with His-GST. In each experiment, “Control” refers to the analyte (either FR or MLCK-FLAG) binding to the ligand-free probe. This control is performed to demonstrate that the association seen in the ligand bound probes is not due to nonspecific binding but is, in fact, specific. The binding of the analyte (either FR or MLCK-FLAG) to the optical probes was measured simultaneously using instrumental defaults for 236 seconds. The dissociation was measured for 287 seconds. There was no binding of FR to the unloaded probes; however, slight binding of MLCK-FLAG to the unloaded probes was observed. The data were analyzed by the Octet data analysis software. The association and dissociation curves for FR binding were globally fit with a 1:1 ligand model, and the curves for MLCK-FLAG binding were locally fit for 80s. The data was plotted using Prism 7.

#### Quantification and Statistical Analysis

##### Analysis of FR-CaM Binding Assay

For CaM^WT^ and CaM^21A,57A^, each data point is presented as the mean ± s.d. of 15 independent experiments. For CaM^94A,130A^, each data point is presented as mean of ± s.d. of 8 independent experiments, and for CaM^21A,57A,94A,130A^, each data point is presented as mean of ± s.d. of 3 independent experiments. For reporter-only data, each data point is presented as a mean ± s.d. of 17 independent experiments. Curve fits were computed using the Matlab (version R2020a) package doseResponse (https://www.mathworks.com/matlabcentral/fileexchange/33604-doseresponse). Algebraic calculations involving the mathematical models were done in Mathematica (version 12.1).

### Statistical Analysis

Statistical calculations were performed using Matlab. AUC was computed using the trapz built-in Matlab function.

#### DNA Sequences

##### CaM Sequences

Red highlighting: mutations

###### His-CaM^WT^ DNA Sequence (protein molecular weight: ∼19.8 kDa)

atgggcagcagccatcatcatcatcatcacagcagcggcctggtgccgcgcggcagccatagcgaaaacctctacttccaatcgatggctgaccagctgactgaggagcagattgcagagttcaaggaggccttctccctctttgacaaggatggagatggcactatcaccaccaaggagttggggacagtgatgagatccctgggacagaaccccactgaagcagagctgcaggatatgatcaatgaggtggatgcagatgggaacgggaccattgacttcccggagttcctgaccatgatggccagaaagatgaaggacacagacagtgaggaggagatccgagaggcgttccgtgtctttgacaaggatgggaatggctacatcagcgccgcagagctgcgtcacgtaatgacgaacctgggggagaagctgaccgatgaggaggtggatgagatgatcagggaggctgacatcgatggagatggccaggtcaattatgaagagtttgtacagatgatgactgcaaagtga

###### His-CaM^21A,57A^ DNA Sequence (protein molecular weight: ∼19.8 kDa)

atgggcagcagccatcatcatcatcatcacagcagcggcctggtgccgcgcggcagccatagcgaaaacctctacttccaatcgatggctgaccagctgactgaggagcagattgcagagttcaaggaggccttctccctctttgccaaggatggagatggcactatcaccaccaaggagttggggacagtgatgagatccctgggacagaaccccactgaagcagagctgcaggatatgatcaatgaggtggcagcagatgggaacgggaccattgacttcccggagttcctgaccatgatggccagaaagatgaaggacacagacagtgaggaggagatccgagaggcgttccgtgtctttgacaaggatgggaatggctacatcagcgccgcagagctgcgtcacgtaatgacgaacctgggggagaagctgaccgatgaggaggtggatgagatgatcagggaggctgacatcgatggagatggccaggtcaattatgaagagtttgtacagatgatgactgcaaagtga

###### His-CaM^94A,130A^ DNA Sequence (protein molecular weight: ∼19.8 kDa)

atgggcagcagccatcatcatcatcatcacagcagcggcctggtgccgcgcggcagccatagcgaaaacctctacttccaatcgatggctgaccagctgactgaggagcagattgcagagttcaaggaggccttctccctctttgacaaggatggagatggcactatcaccaccaaggagttggggacagtgatgagatccctgggacagaaccccactgaagcagagctgcaggatatgatcaatgaggtggatgcagatgggaacgggaccattgacttcccggagttcctgaccatgatggccagaaagatgaaggacacagacagtgaggaggagatccgagaggcgttccgtgtctttgccaaggatgggaatggctacatcagcgccgcagagctgcgtcacgtaatgacgaacctgggggagaagctgaccgatgaggaggtggatgagatgatcagggaggctgccatcgatggagatggccaggtcaattatgaagagtttgtacagatgatgactgcaaagtga

###### His-CaM^21A,57A,94A,130A^ DNA Sequence (protein molecular weight: ∼19.8 kDa)

atgggcagcagccatcatcatcatcatcacagcagcggcctggtgccgcgcggcagccatagcgaaaacctctacttccaatcgatggctgaccagctgactgaggagcagattgcagagttcaaggaggccttctccctctttgccaaggatggagatggcactatcaccaccaaggagttggggacagtgatgagatccctgggacagaaccccactgaagcagagctgcaggatatgatcaatgaggtggcagcagatgggaacgggaccattgacttcccggagttcctgaccatgatggccagaaagatgaaggacacagacagtgaggaggagatccgagaggcgttccgtgtctttgccaaggatgggaatggctacatcagcgccgcag agctgcgtcacgtaatgacgaacctgggggagaagctgaccgatgaggaggtggatgagatgatcagggaggctgccatcgatggagatggccaggtcaattatgaagagtttgtacagatgatgactgcaaagtga

## MLCK FRET REPORTER (FR) DNA Sequence (protein molecular weight: ∼58.2 kDa)

Yellow highlighting: EYFP

Light grey highlighting: Calmodulin binding domain

Blue highlighting: ECFP

Dark grey highlighting: Flexible linker

Green highlighting: FLAG tag

Atggtgagcaagggcgaggagctgttcaccggggtggtgcccatcctggtcgagctggacggcgacgtaaacggccacaagttcagcgtgtccggcgagggcgagggcgatgccacctacggcaagctgaccctgaagttcatctgcaccaccggcaagctgcccgtgccctggcccaccctcgtgaccaccttcggctacggcctgatgtgcttcgcccgctaccccgaccacatgcgccagcacgacttcttcaagtccgccatgcccgaaggctacgtccaggagcgcaccatcttcttcaaggacgacggcaactacaagacccgcgccgaggtgaagttcgagggcgacaccctggtgaaccgcatcgagctgaagggcatcgacttcaaggaggacggcaacatcctggggcacaagctggagtacaactacaacagccacaacgtctatatcatggccgacaagcagaagaacggcatcaaggtgaacttcaagatccgccacaacatcgaggacggcagcgtgcagctcgccgaccactaccagcagaacacccccatcggcaacggccccgtgctgctgcccgacaaccactacctgagctaccagtccgccctgagcaaagaccccaacgagaagcgcgatcacatggtcctgctggagttcgtgaccgccgccgggatcactctcggcatggacgagctgtacaagggtaccgccgctcgtcagaaatggcagaaaaccggacatgcggtgcgtgcgattggccgtctggctgctaccggtagcaagggcgaggagctgttcaccggggtggtgcccatcctggtcgagctggacggcgacgtaaacggccacaggttcagcgtgtccggcgagggcgagggcgatgccacctacggcaagctgaccctgaagttcatctgcaccaccggcaagctgcccgtgccctggcccaccctcgtgaccaccctgacctggggcgtgcagtgcttcagccgctaccccgaccacatgaagcagcacgacttcttcaagtccgccatgcccgaaggctacgtccaggagcgtaccatcttcttcaaggacgacggcaactacaagacccgcgccgaggtgaagttcgagggcgacaccctggtgaaccgcatcgagctgaagggcatcgacttcaaggaggacggcaacatcctggggcacaagctggagtacaactacatcagccacaacgtctatatcaccgccgacaagcagaagaacggcatcaaggcccacttcaagatccgccacaacatcgaggacggcagcgtgcagctcgccgaccactaccagcagaacacccccatcggcgacggccccgtgctgctgcccgacaaccactacctgagcacccagtccgccctgagcaaagaccccaacgagaagcgcgatcacatggtcctgctggagttcgtgaccgccgccgggatcactctcggcatggacgagctgtacaagcccgggggtggatctggtggatctggtggatctatggattacaaggatgacgatgacaag

### Full-length MLCK DNA Sequence (Protein molecular weight: ∼212.0 kDa)

Light grey highlighting: Calmodulin binding domain

Dark grey highlighting: Flexible linker

Green highlighting: FLAG tag

atgggggatgtgaagctggttgcctcgtcacacatttccaaaacctccctcagtgtggatccctcaagagttgactccatgcccctgacagaggcccctgctttcattttgccccctcggaacctctgcatcaaagaaggagccaccgccaagttcgaagggcgggtccggggttacccagagccccaggtgacatggcacagaaacgggcaacccatcaccagcgggggccgcttcctgctggattgcggcatccgggggactttcagccttgtgattcatgctgtccatgaggaggacaggggaaagtatacctgtgaagccaccaatggcagtggtgctcgccaggtgacagtggagttgacagtagaaggaagttttgcgaagcagcttggtcagcctgttgtttccaaaaccttaggggatagattttcagcttcagcagtggagacccgtcctagcatctggggggagtgcccaccaaagtttgctaccaagctgggccgagttgtggtcaaagaaggacagatgggacgattctcctgcaagatcactggccggccccaaccgcaggtcacctggctcaagggaaatgttccactgcagccgagtgcccgtgtgtctgtgtctgagaagaacggcatgcaggttctggaaatccatggagtcaaccaagatgacgtgggagtgtacacgtgcctggtggtgaacgggtcggggaaggcctcgatgtcagctgaactttccatccaaggtttggacagtgccaataggtcatttgtgagagaaacaaaagccaccaattcag atgtcaggaaagaggtgaccaatgtaatctcaaaggagtcgaagctggacagtctggaggctgcagccaaaagcaagaactgctccagcccccagagaggtggctccccaccctgggctgcaaacagccagcctcagcccccaagggagtccaagctggagtcatgcaaggactcgcccagaacggccccgcagactccggtccttcagaagacttccagctccatcaccctgcaggccgcaagagttcagccggaaccaagagcaccaggcctgggggtcctatcaccttctggagaagagaggaagaggccagctcctccccgtccagccaccttccccaccaggcagcctggcctggggagccaagatgttgtgagcaaggctgctaacaggagaatccccatggagggccagagggattcagcattccccaaatttgagagcaagccccaaagccaggaggtcaaggaaaatcaaactgtcaagttcagatgtgaagtttccgggattccaaagcctgaagtggcctggttcctggaaggcacccccgtgaggagacaggaaggcagcattgaggtttatgaagatgctggctcccattacctctgcctgctgaaagcccggaccagggacagtgggacatacagctgcactgcttccaacgcccaaggccaggtgtcctgtagctggaccctccaagtggaaaggcttgccgtgatggaggtggccccctccttctccagtgtcctgaaggactgcgccgttattgagggccaggattttgtgctgcagtgctccgtacgggggaccccagtgccccggatcacttggctgctgaatgggcagcccatccagtacgctcgctccacctgcgaggccggcgtggctgagctccacatccaggatgccctgccggaggaccatggcacctacacctgcctagctgagaatgccttggggcaggtgtcctgcagcgcctgggtcaccgtccatgaaaagaagagtagcaggaagagtgagtaccttctgcctgtggctcccagcaagcccactgcacccatcttcctgcagggcctctctgatctcaaagtcatggatggaagccaggtcactatgactgtccaagtgtcagggaatccaccccctgaagtcatctggctgcacaatgggaatgagatccaagagtcagaggacttccactttgaacagagaggaactcagcacagcctttgtatccaggaagtgttcccggaggacacgggcacgtacacctgcgaggcctggaacagcgctggagaggtccgcacccaggccgtgctcacggtacaagagcctcacgatggcacccagccctggttcatcagtaagcctcgctcagtgacagcctccctgggccagagtgtcctcatctcctgcgccatagctggtgacccctttcctaccgtgcactggctcagagatggcaaagccctctgcaaagacactggccacttcgaggtgcttcagaatgaggacgtgttcaccctggttctaaagaaggtgcagccctggcatgccggccagtatgagatcctgctcaagaaccgggttggcgaatgcagttgccaggtgtcactgatgctacagaacagctctgccagagcccttccacgggggagggagcctgccagctgcgaggacctctgtggtggaggagttggtgctgatggtggtggtagtgaccgctatgggtccctgaggcctggctggccagcaagagggcagggttggctagaggaggaagacggcgaggacgtgcgaggggtgctgaagaggcgcgtggagacgaggcagcacactgaggaggcgatccgccagcaggaggtggagcagctggacttccgagacctcctggggaagaaggtgagtacaaagaccctatcggaagacgacctgaaggagatcccagccgagcagatggatttccgtgccaacctgcagcggcaagtgaagccaaagactgtgtctgaggaagagaggaaggtgcacagcccccagcaggtcgattttcgctctgtcctggccaagaaggggacttccaagacccccgtgcctgagaaggtgccaccgccaaaacctgccaccccggattttcgctcagtgctgggtggcaagaagaaattaccagcagagaatggcagcagcagtgccgagaccctgaatgccaaggcagtggagagttccaagcccctgagcaatgcacagccttcagggcccttgaaacccgtgggcaacgccaagcctgctgagaccctgaagccaatgggcaacgccaagcctgccgagaccctgaagcccatgggcaatgccaagcctgatgagaacctgaaatccgctagcaaagaagaactcaagaaagacgttaagaatgatgtgaactgcaagagaggccatgcagggaccacagataatgaaaagagatcagagagccaggggacagccccagccttcaagcagaagctgcaagatgttcatgtggcagagggcaagaagctgctgctccagtgccaggtgtcttctgaccccccagccaccatcatctggacgctgaacggaaagaccctcaagaccaccaagttcatcatcctctcccaggaaggctcactctgctccgtctccatcgagaaggcactgcctgaggacagaggcttatacaagtgtgtagccaagaatgacgctggccaggcggagtgctcctgccaagtcaccgtggatgatgctccagccagtgagaacaccaaggccccagagatgaaatcccggaggcccaagagctctcttcctcccgtgctaggaactgagagtgatgcgactgtgaaaaagaaacctgcccccaagacacctccgaaggcagcaatgccccctcagatcatccagttccctgaggaccagaaggtacgcgcaggagagtcagtggagctgtttggcaaagtgacaggcactcagcccatcacctgtacctggatgaagttccgaaagcagatccaggaaagcgagcacatgaaggtggagaacagcgagaatggcagcaagctcaccatcctggccgcgcgccaggagcactgcggctgctacacactgctggtggagaacaagctgggcagcaggcaggcccaggtcaacctcactgtcgtggataagccagaccccccagctggcacaccttgtgcctctgacattcggagctcctcactgaccctgtcctggtatggctcctcatatgatgggggcagtgctgtacagtcctacagcatcgagatctgggactcagccaacaagacgtggaaggaactagccacatgccgcagcacctctttcaacgtccaggacctgctgcctgaccacgaatataagttccgtgtacgtgcaatcaacgtgtatggaaccagtgagccaagccaggagtctgaactcacaacggtaggagagaaacctgaagagccgaaggatgaagtggaggtgtcagatgatgatgagaaggagcccgaggttgattaccggacagtgacaatcaatactgaacaaaaagtatctgacttctacgacattgaggagagattaggatctgggaaatttggacaggtctttcgacttgtagaaaagaaaactcgaaaagtctgggcagggaagttcttcaaggcatattcagcaaaagagaaagagaatatccggcaggagattagcatcatgaactgcctccaccaccctaagctggtccagtgtgtgga tgcctttgaagaaaaggccaacatcgtcatggtcctggagatcgtgtcaggaggggagctgtttgagcgcatcattgacgaggactttgagctgacggagcgtgagtgcatcaagtacatgcggcagatctcggagggagtggagtacatccacaagcagggcatcgtgcacctggacctcaagccggagaacatcatgtgtgtcaacaagacgggcaccaggatcaagctcatcgactttggtctggccaggaggctggagaatgcggggtctctgaaggtcctctttggcaccccagaatttgtggctcctgaagtgatcaactatgagcccatcggctacgccacagacatgtggagcatcggggtcatctgctacatcctagtcagtggcctttcccccttcatgggagacaacgataacgaaaccttggccaacgttacctcagccacctgggacttcgacgacgaggcattcgatgagatctccgacgatgccaaggatttcatcagcaatctgctgaagaaagatatgaaaaaccgcctggactgcacgcagtgccttcagcatccatggctaatgaaagataccaagaacatggaggccaagaaactctccaaggaccggatgaagaagtacatggcaagaaggaaatggcagaaaacgggcaatgctgtgagagccattggaagactgtcctctatggcaatgatctcagggctcagtggcaggaaatcctcaacagggtcaccaaccagcccgctcaatgcagaaaaactagaatctgaagaagatgtgtcccaagctttccttgaggctgttgctgaggaaaagcctcatgtaaaaccctatttctctaagaccattcgcgatttagaagttgtggagggaagtgctgctagatttgactgcaagattgaaggatacccagaccccgaggttgtctggttcaaagatgaccagtcaatcagggagtcccgccacttccagatagactacgatgaggacgggaactgctctttaattattagtgatgtttgcggggatgacgatgccaagtacacctgcaaggctgtcaacagtcttggagaagccacctgcacagcagagctcattgtggaaacgatggaggaaggtgaaggggaaggggaagaggaagaagaggtcgacatggactacaaggacgacgacgacaag

## ACKNOWLEDGEMENTS

We would like to thank Dr. Ning Zheng and Dr. Haibin Mao for their help with protein purification, Dr. Martin Golkowski for his help with mass spectrometry, Dr. Anthony Persechini for providing the MLCK FR construct, and Dr. Zenon Grabarek and Dr. John Biddle for comments on the manuscript. The authors are also grateful to UW’s elevator maintenance team for rescuing MJSM (and several liters of induced bacteria) from a stalled elevator during the writing of this manuscript.

## AUTHOR CONTRIBUTIONS

MJSM, JPD, and YS designed the experiments and wrote the manuscript. MJSM, HE, and YS performed cloning, binding assays, and other biochemical experiments. DVR performed BLI experiments. TL and HG contributed to experimental design. MJSM, JPD, and YS analyzed the data. JPD developed the mathematical models.

## FUNDING AND ADDITIONAL INFORMATION

MJSM was supported by NIH grant T32GM007750; JPD was supported by a Neukom Fellowship and a Harvard Data Science Fellowship.

## DATA AVAILABILITY

All raw data used for plate reader experiments and calculations are available upon request.

## ABBREVIATIONS AND NOMENCLATURE

CaM^WT^: – wild-type calmodulin
CaM^21A,57A^: – calmodulin with N-terminus D21A and D57A mutations
CaM^94A,130A^: – calmodulin with C-terminus D94A and D130A mutations
CaM^21A,57A,94A,130A^: – calmodulin with N-terminus D21A and D57A mutations, and C-terminus D94A and D130A mutations
MLCK-FLAG: – full-length MLCK protein tagged with FLAG
FR: – FRET Reporter
FRET: – fluorescence resonance energy transfer
HEK 293T: – human embryonic kidney 293T cells
MLCK: – smooth muscle or nonmuscle myosin light chain kinase
STR analysis: – short tandem repeat analysis

---

**Supplementary Figure 1:**
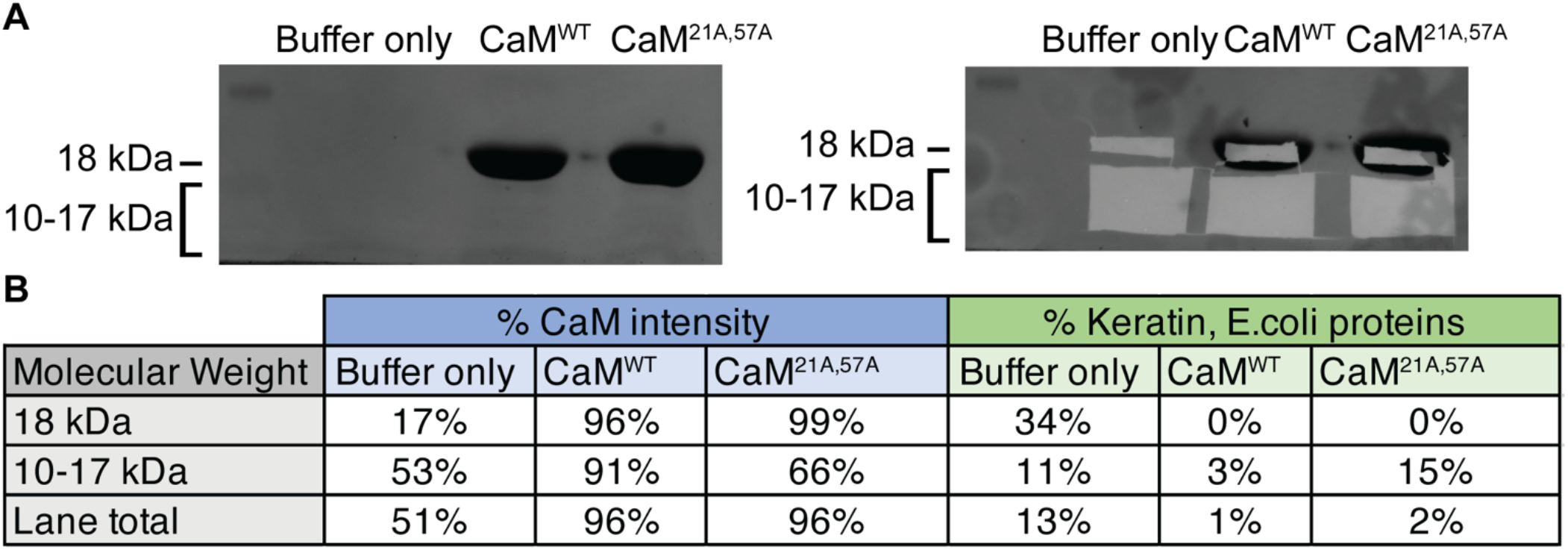
Mass spectrometry analysis of purified calmodulin proteins. Semi-quantitative mass spectrometry was performed to assess the purity of purified CaM^WT^ and CaM^21A,57A^ proteins; a “loading buffer only” lane was used as a control. **(A)** Following SDS-PAGE, the gel was Coomassie stained, and bands of interest were excised and analyzed by mass spectrometry. **(B**) The identified peptides were categorized as CaM or a contaminating protein (keratin or *E. coli* protein) using their annotations. Finally, the percent abundance of either CaM or contaminants in each band was determined using their intensity in mass spectrometry data

**Supplementary Figure 2:**
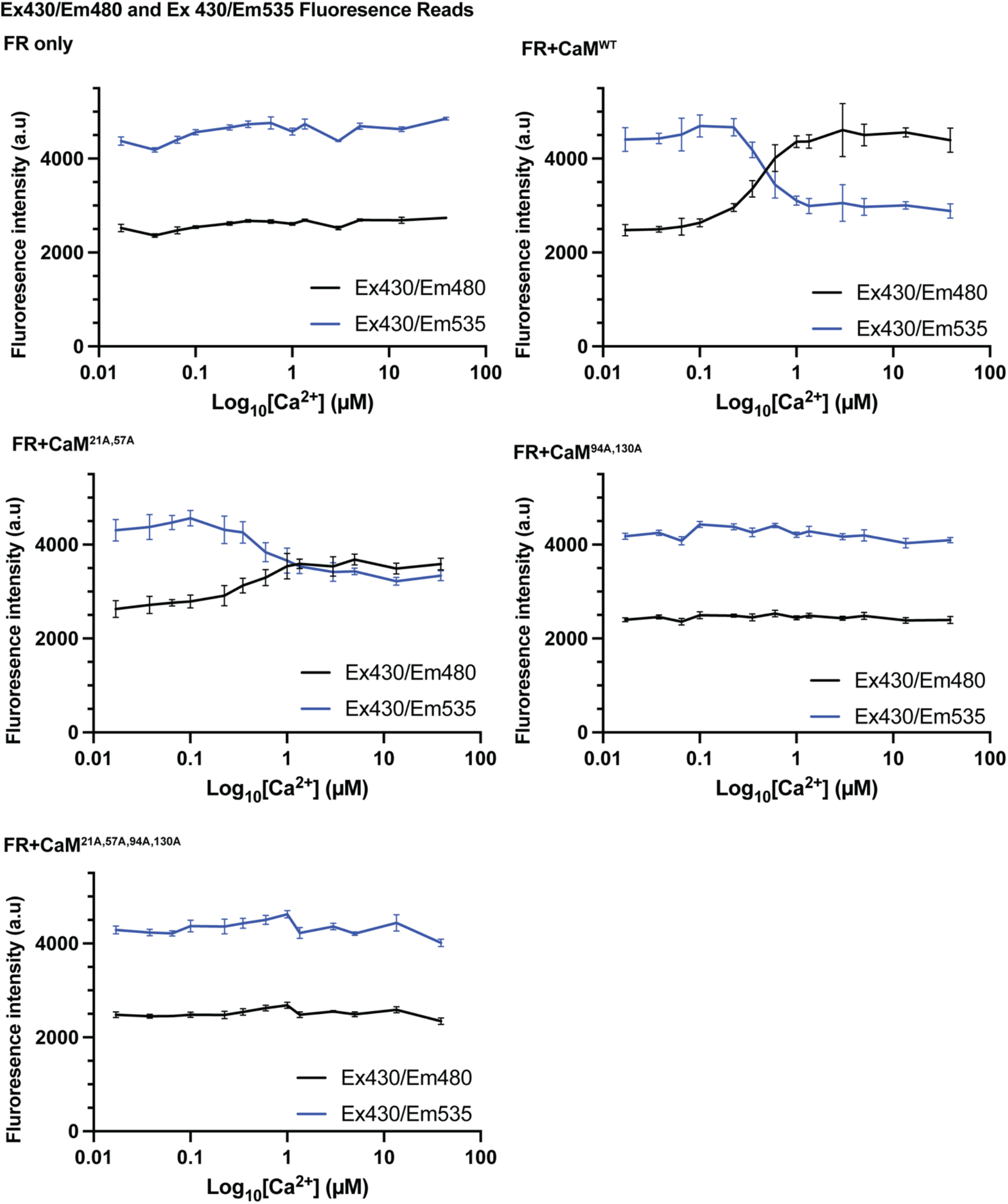
Fluorescence intensities of FR-CaM binding assay. For each of the indicated CaM-FR combinations, 673 nM CaM and 22.9 nM FR were combined in 150 μl of Ca^2+^ buffer (14 conditions between 0 μM and 39 μM). Fluorescence intensity of [Ex 430, Em 480], followed by intensity of [Ex 430, Em 535], was measured using a plate reader. Shown are mean ± standard deviation of raw data from at least 5 replicates for CaM^WT^, CaM^21A,57A^, and CaM^94A,130A^, and 3 replicates for CaM^21A,57A,94A,130A^ and FR only. **Fig. 3** shows the full dataset following analysis.

**Supplementary Figure 3:**
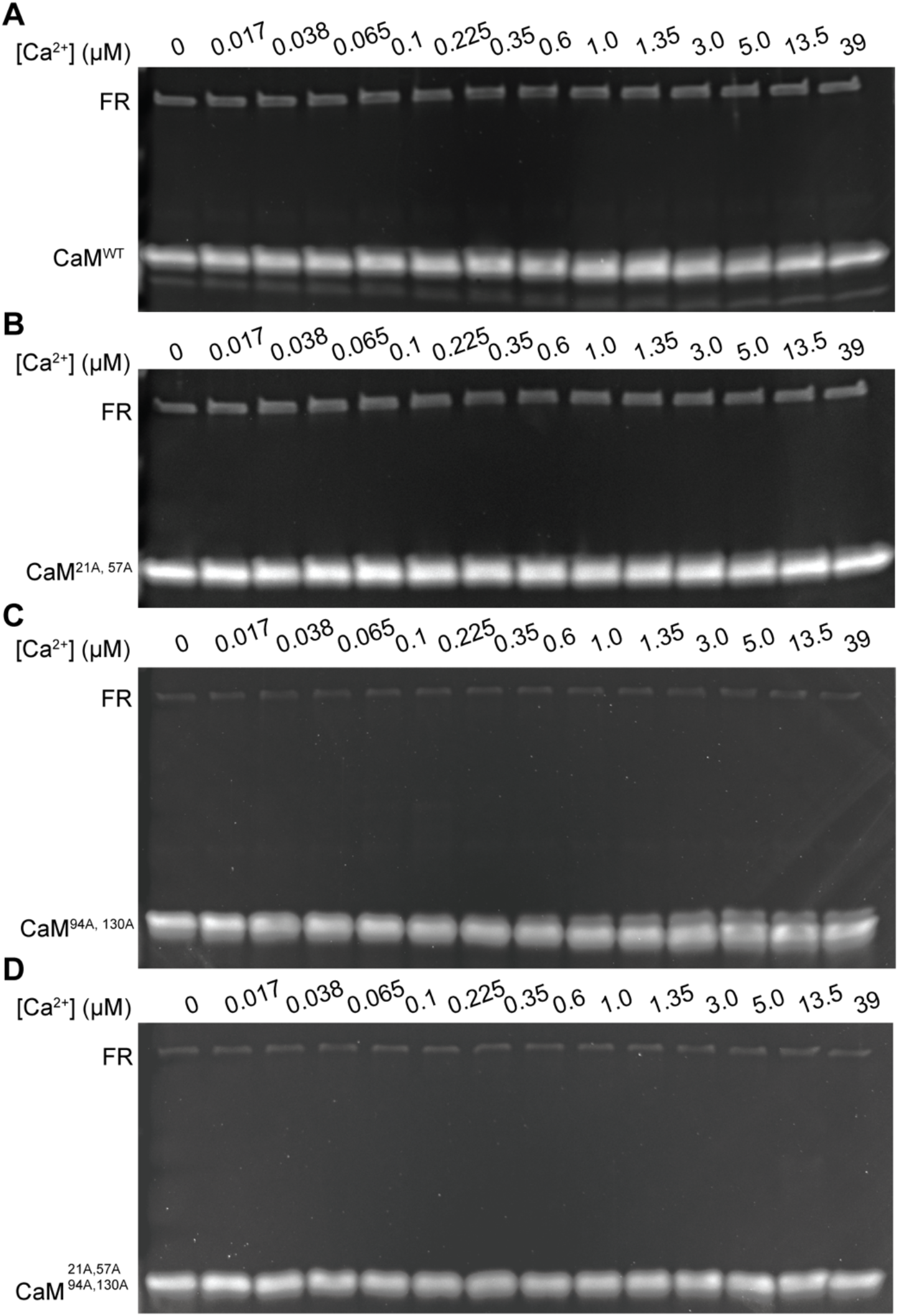
Purified FR and CaM used in data collection. A portion of each sample for fluorescent detection of FR-CaM binding was recovered from the wells. SDS-PAGE and SYPRO™ Ruby staining confirmed the presence of excess CaM compared to FR in the FRET-based binding assay for **(A)** CaM^WT^ and FR, **(B)** CaM^21A,57A^ and FR, **(C)** CaM^94A,130A^ and FR, and **(D)** CaM^21A,57A,94A,130A^

**Supplementary Figure 4.**
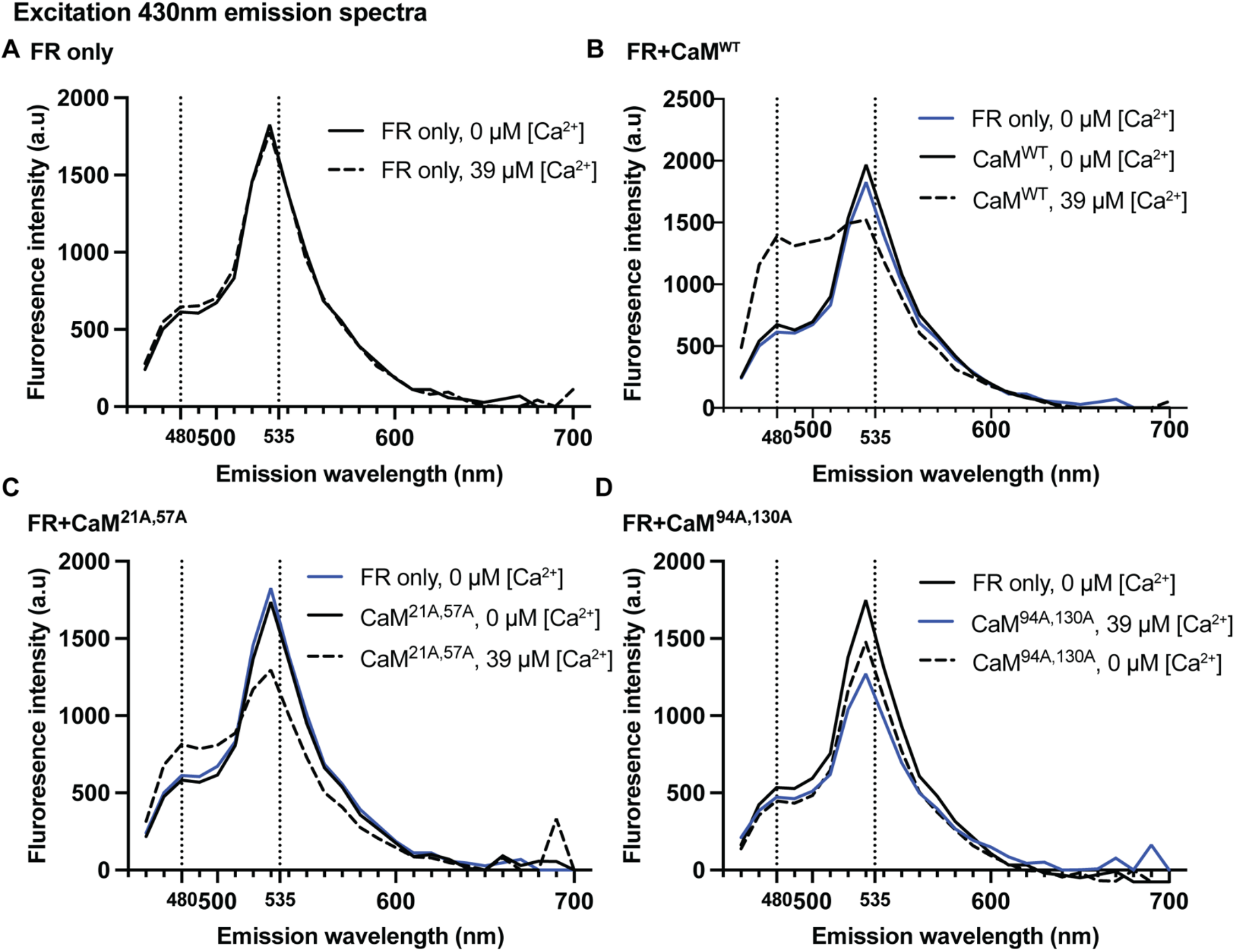
Emission spectra of FR at Ex 430. The emission spectra (460 nm - 700 nm, 10 nm steps) of **(A)** FR alone or with **(B)** CaM^WT^, **(C)** CaM^21A,57A^, or **(D)** CaM^94A,130A^ was measured at either 0 or 39 μM [Ca^2+^]. Dotted lines indicate the Em 480 and Em 535 wavelengths, which were used in FRET-based FR-CaM binding assays. One representative read for each condition is shown. FR-only spectra are included in the CaM graphs as a reference.

**Supplementary Figure 5:**
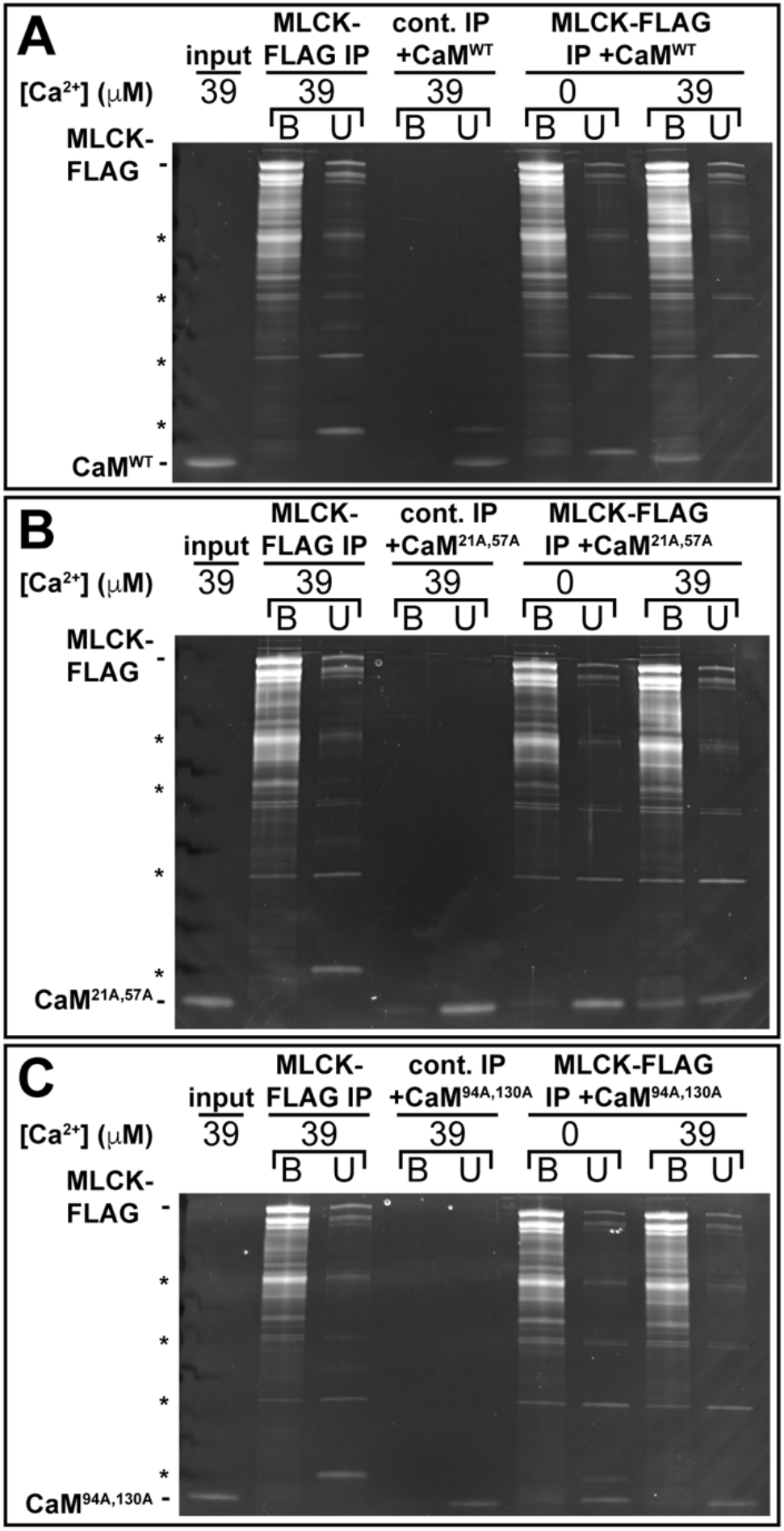
Purified CaM^WT^ and CaM^21A,57A^ show Ca^2+^-dependent interaction with MLCK-FLAG; CaM^94A,130A^ does not. Representative images showing binding between bead-immobilized MLCK-FLAG and **(A)** CaM^WT^, **(B)** CaM^21A,57A^, and **(C)** CaM^94A,130A^ in buffers with the indicated free [Ca^2+^]. Unbound and bound fractions were analyzed by SDS-PAGE followed by SYPRO™ Ruby protein gel stain. MLCK-FLAG without CaM addition and a control binding experiment using non-transfected HEK 293T cells serve as controls. The input lane shows the purified CaM^WT^, CaM^21A,57A^, or CaM^94A,130A^ added to the on-bead binding assay. Asterisks (*) indicates background bands.

**Supplementary Figure 6.**
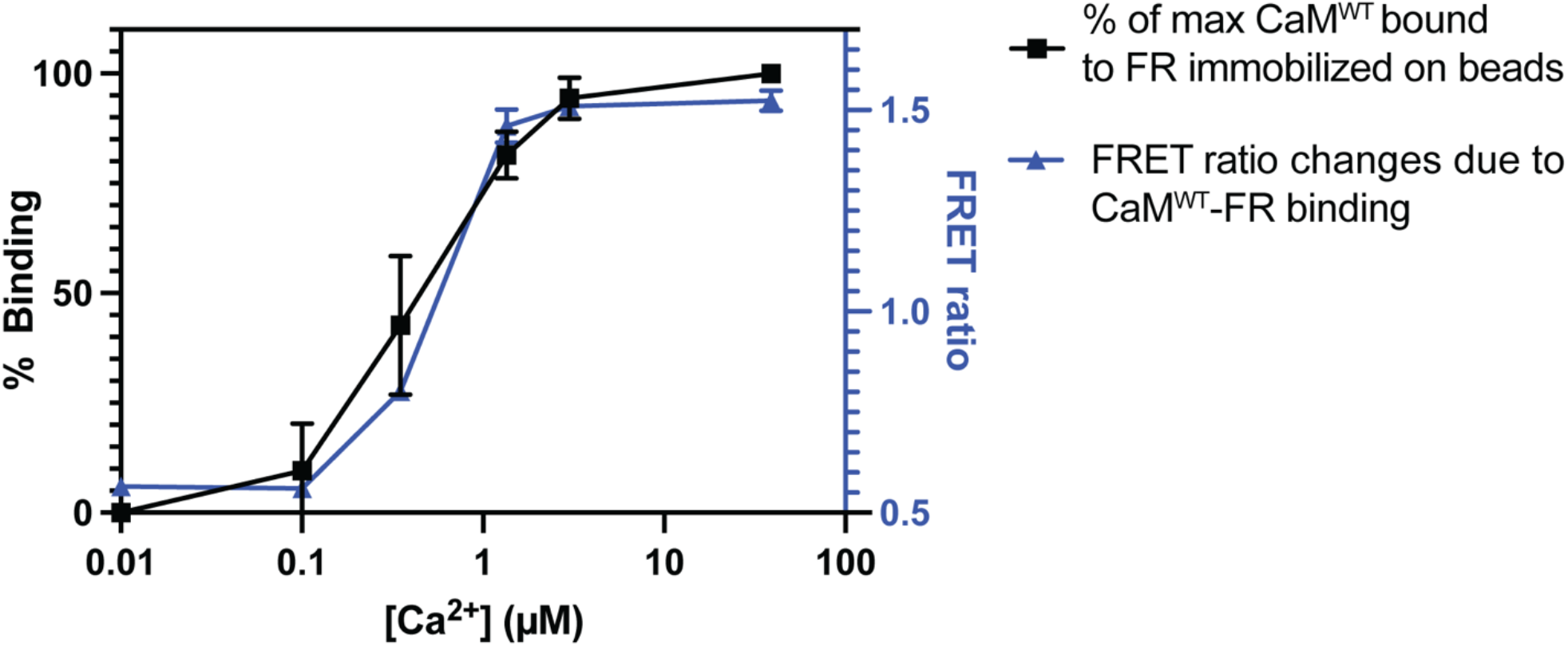
Comparison of on-bead and FRET-based binding assessment between FR and CaM^WT^ as a function of [Ca^2+^]. Data collected using FRET-based binding assay for CaM^WT^-FR was plotted alongside data collected using on-bead binding assay. To quantify the relative amount of CaM bound to FR, CaM^WT^-FR binding at 39 μM [Ca^2+^] was set to 100%, binding at other [Ca^2+^] relative to this maximum signal were calculated.

**Supplementary Figure 7:**
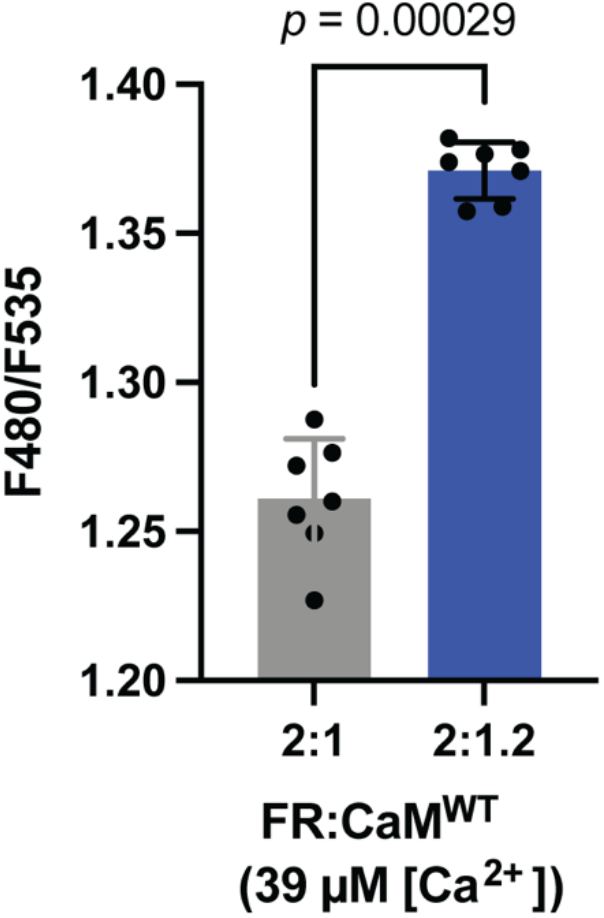
FR-CaM binding assay is sensitive enough to detect a calculated 8.5% increase in CaM-FR binding. A modified version of the FR-CaM FRET-based binding assay using a FR:CaM^WT^ ratio of either 2:1 or 2:1.2 (22.9 nM FR and 11.45 nM or 13.75 nM CaM) in high [Ca^2+^] was performed. The Em 480/Em F535 ratio increased significantly between conditions (*p* = 0.00029 by a one-tailed Mann-Whitney *U* test), from an average of 1.26 to 1.37 (an 8.7% increase in FRET ratio). Assuming Model 2 and the reference parameter values from Fajmut, Brumen *et al*. (*9*), the fraction of FR bound to CaM^WT^ is predicted to differ by 8.5% between the two conditions (46.2% vs. 54.7% bound). Mean ± standard deviation of 7 replicates for each condition are shown.

**Supplementary Figure 8:**
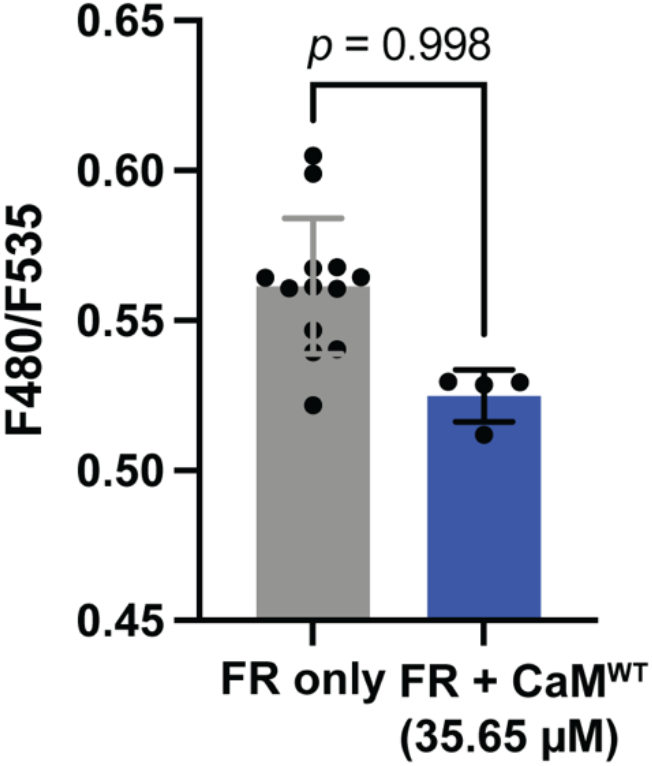
CaM^WT^ does not bind to FR in the absence of Ca^2+^ even at high concentrations. To test the robustness of our observation of no binding in zero [Ca^2+^], we repeated the FRET-based binding assay with 35.65 μM CaM^WT^, for which Model 1 predicts 21.7% binding even if previous estimates of a key parameter value are incorrect by an order of magnitude. CaM^WT^ was concentrated to 66.6 μg/μl so that the volume of protein solution used was comparable to previous assays. As in the original assay, the F480/F535 ratio was not higher than baseline (*p* = 0.998 by a one-tailed Mann-Whitney *U* test). Mean ± standard deviation of 13 replicates for FR only read and 4 replicates for FR+ CaM^WT^ reads are shown.

**Supplementary Figure 9.**
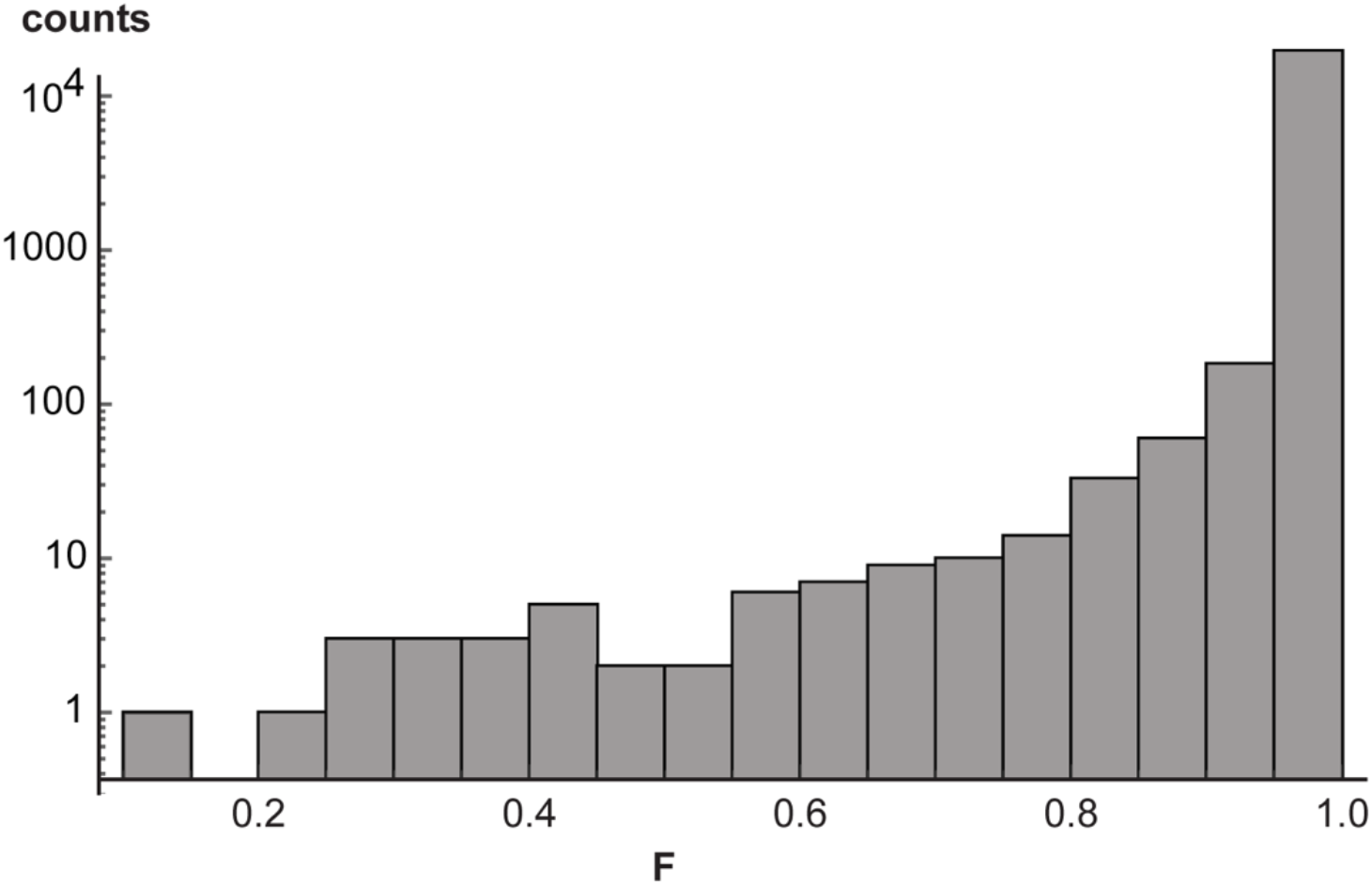
Histogram from sensitivity analysis of model of CaM^21A,57A^. The histogram shows the distribution of predicted binding fractions for 200,000 combinations of K_4_ and K_6_ chosen at random from the interval [0.1v, 10v], where v is the reference value from Fajmut, Brumen *et al*. (*9*).

**Supplementary Figure 10.**
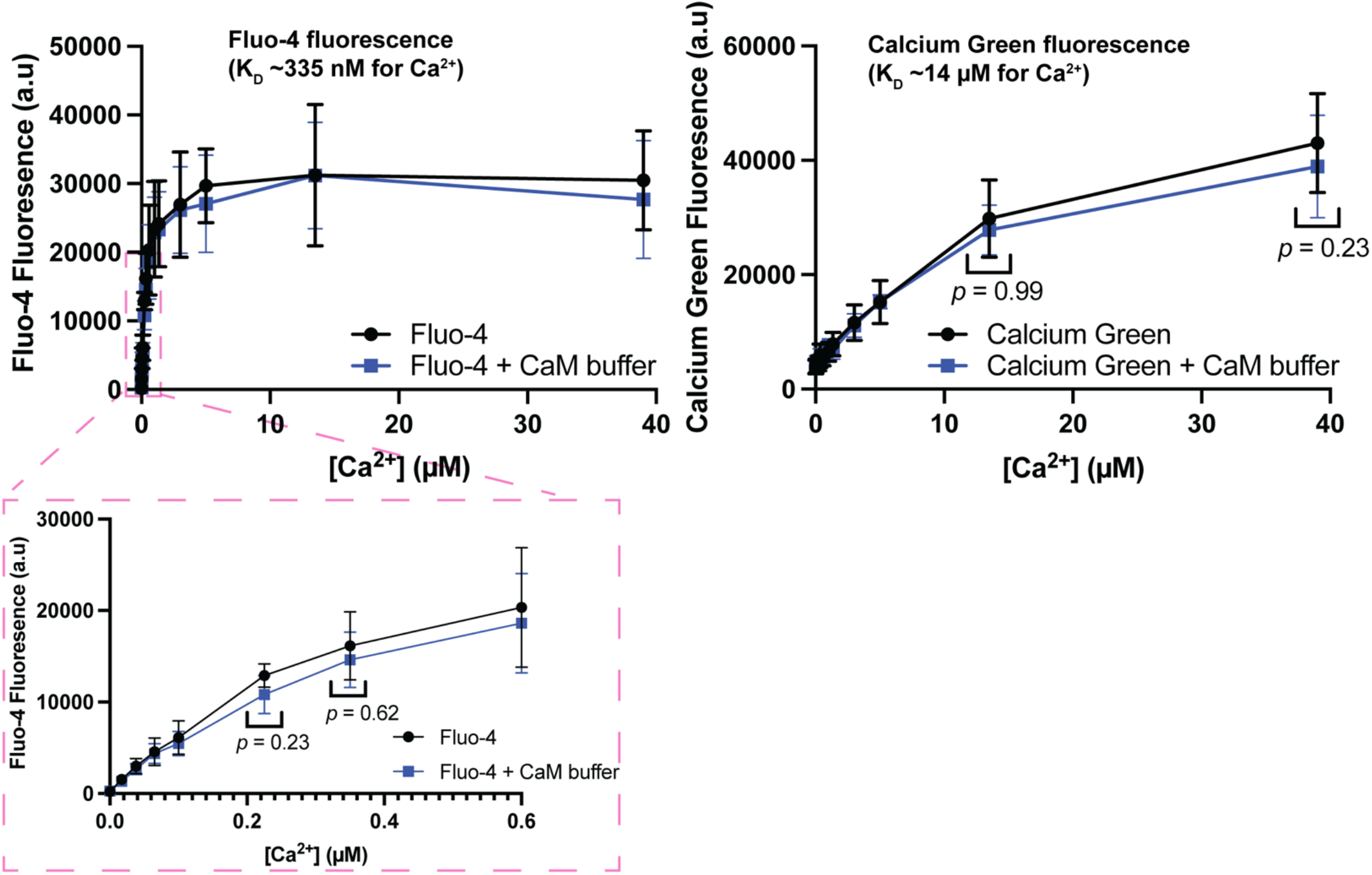
Buffer addition to FR-CaM binding assay does not change assay free [Ca^2+^]. Ca^2+^ indicator dyes Fluo-4 or Calcium Green were added to the 14-buffer series of the FR-CaM binding assay; data was collected using a plate reader with a GFP filter. The same buffer/Ca^2+^ indicator dye combinations were then assayed following the addition of a “dummy buffer” composed of the buffer in which all CaM proteins were stored. The area of interest (magnified region in red dotted line) highlights the data corresponding to [Ca^2+^] most relevant to Fluo-4, which has K_D_∼335 nM. Calcium Green has K_D_∼14 μM. Mean ± standard deviation of four replicates for each condition are shown. *P* values were calculated using a two-tailed Mann-Whitney *U* test.

